# Lack of phosphatidylinositol 3-kinase VPS34 in regulatory T cells leads to a fatal lymphoproliferative disorder without affecting their development

**DOI:** 10.1101/2024.01.08.574346

**Authors:** Christina J.F. Courreges, Elizabeth C.M. Davenport, Benoit Bilanges, Elena Rebollo-Gomez, Jens Hukelmann, Priya Schoenfelder, James R. Edgar, David Sansom, Cheryl Scudamore, Rahul Roychudhuri, Oliver A. Garden, Bart Vanhaesebroeck, Klaus Okkenhaug

## Abstract

Regulatory T (Treg) cells are essential for the maintenance of immunological tolerance, yet the molecular components required for their maintenance and effector functions remain incompletely defined. Inactivation of VPS34 in Treg cells led to an early, lethal phenotype, with massive effector T cell activation and inflammation, like mice lacking Treg cells completely. However, VPS34-deficient Treg cells developed normally, populated the peripheral lymphoid organs and effectively supressed conventional T cells *in vitro*.

Our data suggest that VPS34 is required for the maturation of Treg cells or that mature Treg cells depend on VPS34 for survival. Functionally, we observed that lack of VPS34 activity impairs cargo processing upon transendocytosis, that defective autophagy contributes to, but is not sufficient to explain this lethal phenotype, and that loss of VPS34 activity induces a state of heightened metabolic activity that may interfere with metabolic networks required for maintenance or suppressive functions of Treg cells.

## Introduction

Immune suppression by regulatory T cells (Treg cells) is essential for immunological tolerance, the mechanism through which the immune system is restrained from mounting an attack on self-tissues, commensal organism and innocuous foreign antigens (i.e. allergens) (Izcue et al., 2009; Josefowicz et al., 2012; Wing and Sakaguchi, 2010). Indeed, the absence of Treg cells, caused by loss-of-function mutations in their defining transcription factor *Foxp3,* leads to lethal autoimmunity, both in humans and mice (Brunkow et al., 2001; Fontenot et al., 2017; Wildin et al., 2001).

Treg cells are characterised by the constitutive expression of the interleukin-2 (IL-2) receptor and cytotoxic T-lymphocyte-associated protein 4 (CTLA-4). CTLA-4 on Treg cells can capture CD80 and CD86 from antigen-presenting cells through a process termed transendocytosis, thus depriving conventional T cells (Tcon) of these costimulatory ligands (Omar S Qureshi et al., 2011). Similarly, internalisation and degradation of extracellular IL-2 by the IL-2 receptor on Treg cells leads to depletion of this cytokine from the cellular environment, thereby dampening Tcon activation (Pandiyan et al., 2007). Treg cells also secrete immunosuppressive cytokines such as IL-10 (Asseman et al., 1999) and transforming growth factor (TGF)-β (Fahlén et al., 2005), as well as inhibitory metabolites (Deaglio et al., 2007). However, our understanding of the suppressive mechanisms used by Treg cells to maintain immunological tolerance remains incomplete.

The PI3K VPS34 phosphorylates the phosphatidylinositol (PI) lipid on the 3-position of the inositol ring to generate phosphatidylinositol 3-phosphate (PI3P) (Backer, 2016). VPS34 forms two distinct protein complexes that control different cellular processes: VPS34 complex 1 is associated with autophagy, whereas VPS34 complex 2 controls endosomal trafficking. Both endosomal traffic and autophagy are processes known to be important for Treg cell function. Indeed, CTLA-4 and the IL-2 receptor-dependent mechanisms of Treg cell suppression depend on endocytosis. Autophagy is critical to maintain cellular homeostasis of immune cells and is required for normal Treg cell function (Kabat et al., 2016; Wei et al., 2016).

Previous studies showed that deletion of *Pik3c3*, the gene encoding VPS34, resulted in defective T cell homeostasis. This was either ascribed to impaired IL-7 receptor recycling (McLeod et al., 2011) or impaired processing of autophagosomes (Parekh et al., 2013; Willinger and Flavell, 2012). Deletion of *Pik3c3* in T cells also led to the development of an inflammatory disease in older mice that was correlated with a loss of Treg cell homeostasis (Parekh et al., 2013). Yang et *al*. reported that VPS34 deletion in T cells led to reduced mitochondrial membrane potential and impaired oxidative phosphorylation (OXPHOS) after T cell activation (Yang et al., 2020). Nevertheless, the specific role of VPS34 in Treg cells remains unknown and is mostly inferred from studies of other cell types and non-mammalian organisms. Therefore, we explored whether VPS34 is critical for Treg cell-mediated suppression using a conditional knock-out mouse model. We found that mice with VPS34 inactivated in Treg cells died within 6 weeks of birth from an autoimmune lymphoproliferative disease, similar to what has previously been observed in *Foxp3* knockout and *Scurfy* mice (Brunkow et al., 2001). However, in contrast to *Foxp3* knockout mice, and most other models that recapitulate the *Scurfy* phenotype, VPS34-deficient Treg cells develop normally and populate the peripheral lymphoid organs. Moreover, VPS34-deficient Treg cells could suppress Tcon *in vitro*. However, when investigating the transendocytosis of CD80 via CTLA-4 and the subsequent degradation of the cargo, inhibition of VPS34 led to impaired clearance of the endocytosed material. Proteomic profiling provided indications that VPS34 inhibition alters metabolic functions of Treg cells, possibly as a failure to effectively clear mitochondria. Together, this may lead to fragility especially of activated Treg cells and consequently loss of Treg cell–mediated immunological tolerance.

## Results

### Deletion of VPS34 in Treg cells leads to a Scurfy-like phenotype

To create a conditional VPS34 allele, we inserted *loxP* sites flanking exon 21 of *Pik3c3*, which encodes a critical stretch of 25 amino acids (Ala730 to Thr754) in the VPS34 kinase domain, producing an in-frame deletion upon Cre-mediated recombination (**Fig. S1A, B)**. In line with our expectations, overexpression of a c-Myc tagged construct of this deletion mutant showed a lower molecular weight band than for the c-Myc tagged WT construct (**Fig. 1A**), confirming that the deletion of 25 amino acid maintains the open reading frame, thus creating a truncated version of VPS34. Importantly, this deletion renders the truncated VPS34 protein catalytically inactive (**Fig. 1A**).

**Fig. 1.**
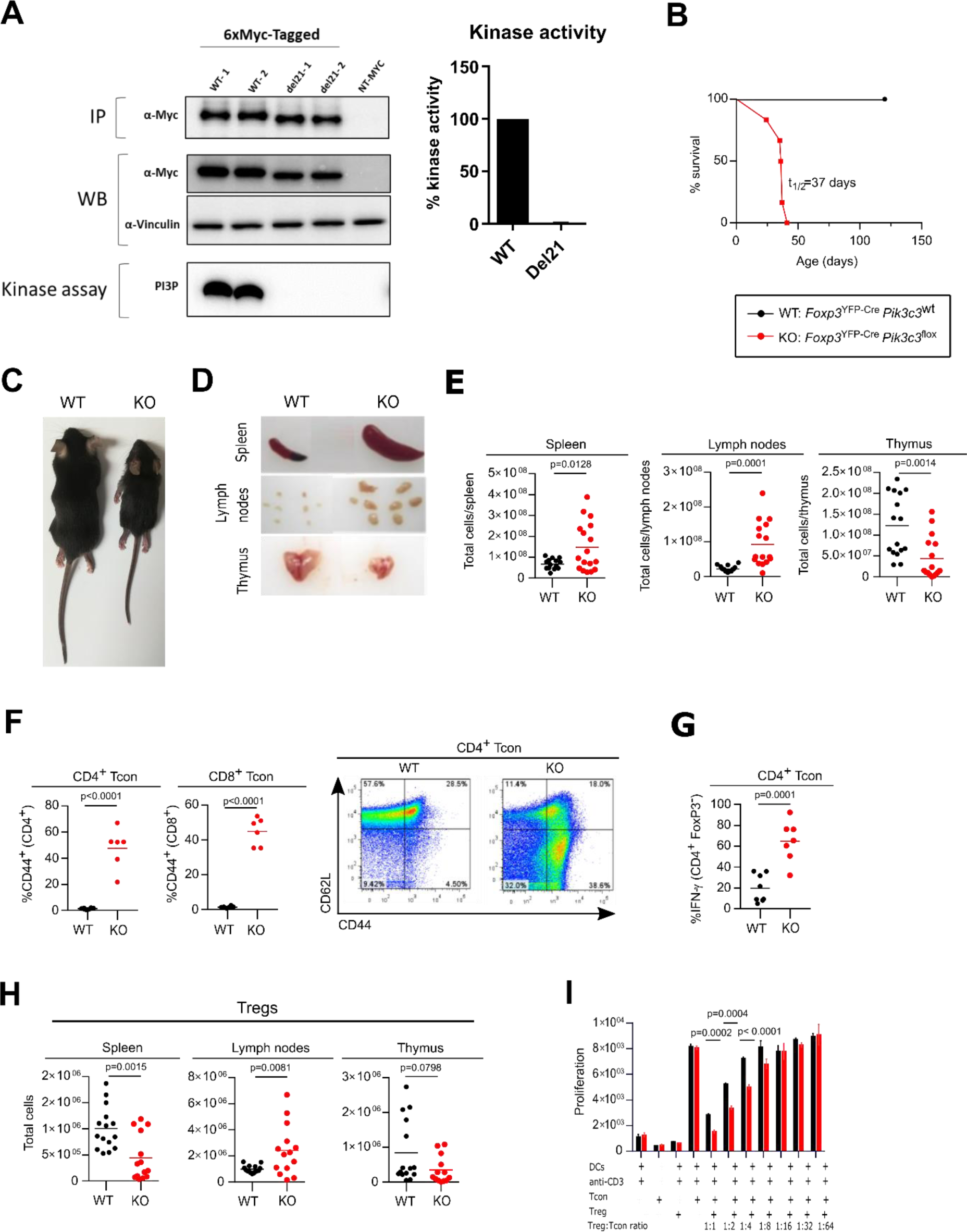
Deletion of VPS34 in Treg cells leads to a Scurfy-like phenotype. **A)** Immunoprecipitation (IP), expression (WB) and *in vitro* kinase assay of tagged versions of the truncated (del21) and wild-type (WT) allele in transiently-transfected HEK293 cells. IPs were performed using anti-Myc antibodies. NT = non-transfected controls. *In vitro* lipid kinase assay was performed using PI as a substrate on 6xMyc-tagged VPS34 IPs. Representative data of 3 independent experiments is shown with 2 replicates per experiments (labelled as WT-1, WT-2, del21-1, del21-2). **B)** Survival curves for *Foxp3*^YFP-Cre^*Pik3c3*^flox^ (red) and wild-type *Foxp3*^YFP-Cre^*Pik3c3*^WT^ mice (black). **C)** Representative pictures of wild-type *Foxp3*^YFP-Cre^ *Pik3c3*^WT^ mice (left) and Foxp*3*^YFP-Cre^*Pik3c3*^flox^ (right). **D**) Examples of enlarged spleen and lymph nodes from *Foxp3*^YFP-Cre^*Pik3c3*^flox^ mice and wild-type (WT) littermate controls. **E**) Absolute numbers of cells in the spleen, lymph nodes (inguinal, brachial, and axillary) and thymus from *Foxp3*^YFP-Cre^*Pik3c3*^flox^ mice compared to wild-type mice. **F**) Percentage and representative FACS plots of CD44^high^ CD62^low^ Tcon and CD8^+^ T cells in the spleen. **G**) Percentage of IFN-γ^+^ CD4^+^ CD25^-^ T cells in the spleen. **H**) Absolute numbers of Treg cells in spleen, lymph nodes (inguinal, brachial, and axillary), and thymus of Foxp*3*^YFP-Cre^*Pik3c3*^flox^ and wild-type *Foxp3*^YFP-Cre^ *Pik3c3*^WT^ mice. **I**) Tcon and Treg cells from *Foxp3*^YFP-Cre^*Pik3c3*^flox^ mice and control wild-type mice (WT) were co-cultured in the presence of dendritic cells and 1μg/ml anti-CD3 for 96 h. Wells were pulsed with Alamar Blue for 4 h and the measured fluorescence used to gauge proliferation. Error bars represent SEM of triplicate wells. *Foxp3*^YFP-Cre^*Pik3c3*^flox^ mice and the respective control mice were between 4 and 5.5 weeks of age. *Foxp3*^YFP-Cre/WT^*Pik3c3*^flox^ mosaic mice and the respective control mice were between 8 and 12 weeks of age. n = 3-15 mice per group. Statistical significance was determined using an unpaired two-tailed Student’s t-test (c, d, e, i, h), paired two-tailed Student’s t-test (**A, B, F, G, J**), or a One-way ANOVA with Tukey’s post-test (**I**). Results are pooled from 2 to 4 independent experiments

Crosses of *Pik3c3*^flox^ mice with Cre-deleter (B6.C-Tg(CMV-Cre )1Cgn/J) transgenic mice (which ubiquitously express the Cre recombinase from the zygote stage of development (Schwenk et al., 1995)) showed that neither homozygous VPS34^Del21/Del21^ mutant embryos at E9.5d or pups were viable, confirming the importance of VPS34 in embryonic development (data not shown). Altogether, this confirms that truncation of this critical stretch in the VPS34 kinase domain recapitulates the embryonic lethality previously reported using different *Pik3c3* gene targeting strategies (Bilanges et al., 2017; Zhou et al., 2011).

To explore whether VPS34 is critical for Treg cell-mediated suppression, we generated a conditional knockout (KO) mouse by crossing *Pik3c3*^flox^ with *Foxp3*^YFP-Cre^ mice (**Fig. S1C, D**). Remarkably, mice with VPS34 inactivated in Treg cells exhibited smaller body size, hunched posture, and crusting and scaling of the ears and abdomen, and became moribound within 4-6 weeks of birth (t_1/2_: 37 days) (**Fig. 1B, C**). *Foxp3*^YFP-Cre^*Pik3c3*^flox^ mice also developed splenomegaly and lymphadenopathy (**Fig. 1D**), suggesting an ongoing autoimmune lymphoproliferative disease, similar to what has previously been observed in *Foxp3* knockout and Scurfy mice (Brunkow et al., 2001). By contrast, the thymus of *Foxp3*^YFP-Cre^*Pik3c3*^flox^ mice was reduced in size (**Fig. 1D**), which is likely secondary to the ongoing inflammation. Accordingly, total cell numbers were increased in the spleen and lymph nodes while reduced in the thymus (**Fig. 1E**). Histopathological review revealed marked infiltration of lymphocytes, macrophages, and neutrophils into secondary lymphoid organs, i.e. the spleen and lymph nodes, as well as multiple organs such as liver, lung and bone marrow (**Fig. S1E**). In line with the ongoing lymphoproliferative disease, *Foxp3*^YFP-Cre^*Pik3c3*^flox^ mice had substantial expansion of CD4^+^ and CD8^+^ T cells (**Fig. S1F and G, respectively**) in the lymph nodes and the spleen, and the majority of CD4^+^ and CD8^+^ T cells in *Foxp3*^YFP-Cre^*Pik3c3*^flox^ mice expressed the activation marker CD44. Additionally, CD4^+^ T cells produced increased levels of IFN-γ, suggesting systemic hyperactivation of the T cell compartment (**Fig. 1F, G**). *Foxp3*^YFP-Cre^*Pik3c3*^flox^ mice had increased numbers of Treg cells in their lymph nodes, but fewer in the spleen, and normal numbers in the thymus (**Fig. 1H**). However, the proportions of Treg cells were reduced in the lymph nodes and the spleens of *Foxp3*^YFP-Cre^*Pik3c3*^flox^ mice, while increased in the thymi (**Fig. S1H**). The expansion of activated T cells in the presence of Treg cells suggests that the loss of VPS34 interferes with Treg cell suppressive functions rather than simply by preventing their development.

Surprisingly, VPS34-deficient Treg cells from *Foxp3*^YFP-Cre^*Pik3c3*^flox^ mice supressed the proliferation of Tcon *in vitro* more efficiently than their wild-type counterparts (**Fig. 1I**). VPS34-deficient Treg cells also caused similar dose-dependent reduction of the IL-2 concentration in the co-culture assay medium (**Fig. S1I**). These results demonstrate that Treg cells that develop in the absence of VPS34 do maintain functions associated with WT Treg cells *in vitro*, yet fail to prevent lethal autoimmune disease *in vivo*.

### VPS34-deficient Treg cells have a competitive disadvantage, but are not intrinsically pathological

As *Foxp3*^YFP-Cre^*Pik3c3*^flox^ mice developed such a profound inflammatory disease, we sought to determine whether the observed differences between VPS34-sufficient and VPS34-deficient Treg cells were due to an intrinsic role of VPS34 in Treg cells or whether the altered Treg cell phenotype was secondary to the inflammatory milieu. We therefore generated mosaic knockout mice by taking advantage of the localisation of the *Foxp3* gene on the X chromosome (**Fig. S2A**). Female mice heterozygous for the *Foxp3*^YFP-Cre^ transgene (*Foxp3*^YFP-Cre/WT^) should delete VPS34 in approximately 50% of their Treg cells after random X chromosome inactivation (I Okamoto, A P Otte, C D Allis, D Reinberg, 2004). *Foxp3*^YFP-Cre/WT^*Pik3c3*^flox^ mosaic mice were healthy (data not shown), demonstrating that the presence of functional Treg cells can prevent the development of a fatal lymphoproliferative disease even when VPS34 is deleted in about half of the total Treg cells. This argues against a dominant pathological role of VPS34-deficient Treg cells.

*Foxp3*^YFP-Cre/WT^*Pik3c3*^flox^ mosaic mice had normal CD4^+^ and CD8^+^ T cell numbers (**Fig. S2B, C**, respectively), which displayed a normal level of the activation marker CD44 (**Fig. S2D**). Similarly, the proportions (**Fig. 2A**) and cellularity (**Fig. S2E**) of Treg cells in the spleens, lymph nodes and thymi of *Foxp3*^YFP-Cre/WT^*Pik3c3*^flox^ mosaic mice were comparable to those of control mice. However, the proportion of VPS34-deficient Treg cells (YFP^+^ Cre^+^ Treg cells from *Foxp3*^YFP-Cre/WT^*Pik3c3*^flox^ mosaic mice) was reduced compared to the proportion of VPS34-sufficient Treg cells (YFP^+^ Cre^+^ Treg cells from *Foxp3*^YFP-Cre/WT^*Pik3c3*^WT^ control mice) (**Fig. 2B**). These data indicate that while VPS34-deficient Treg cells are not intrinsically pathological, they have a competitive disadvantage compared to VPS34-sufficient Treg cells.

**Fig. 2.**
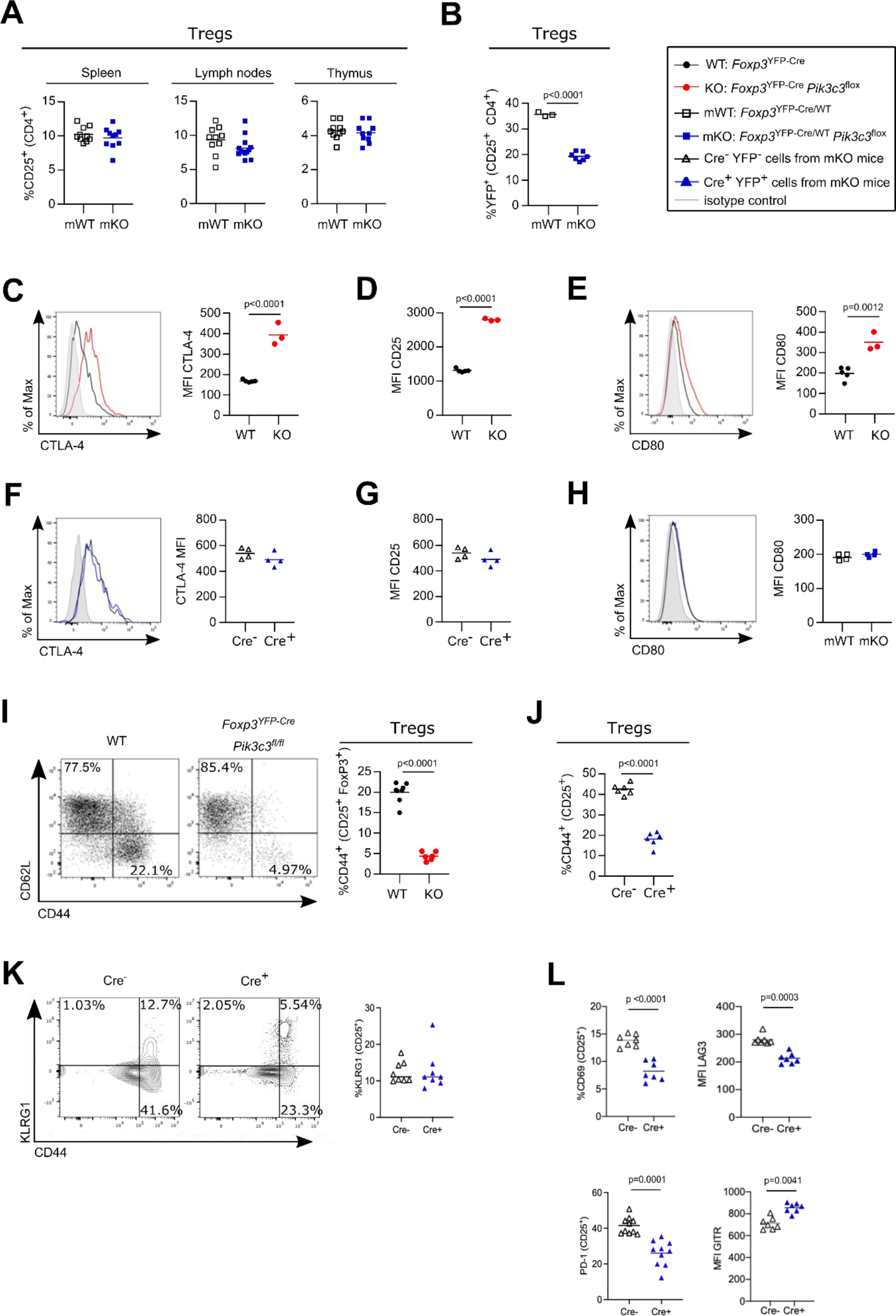
VPS34-deficient Treg cells are not intrinsically pathological, but have a competitive disadvantage. **A)** Percentage of CD25^high^ cells from CD4^+^ in the spleen, the lymph nodes (inguinal, brachial, and axillary), and the thymus of *Foxp3*^YFP-Cre/WT^*Pik3c3*^flox^ mosaic mice and *Foxp3*^YFP-Cre/WT^ control mice (WT). **B**) Percentage of YFP^+^ cells among CD25^+^ CD4^+^ cells in the lymph nodes (inguinal, brachial, and axillary) of *Foxp3*^YFP-Cre/WT^*Pik3c3*^flox^ mosaic mice and *Foxp3*^YFP-Cre/WT^ *Pik3c3*^WT^ control mice (WT). **C – D**) Mean fluorescence intensity (MFI) of CTLA-4 (**C**) and CD25 (**D**) on splenic Treg cells from *Foxp3*^YFP-Cre^*Pik3c3*^flox^ mice and *Foxp3*^YFP-Cre^ control mice (WT). **E – F**) Mean fluorescence intensity (MFI) of CTLA-4 (**E**) and CD25 (**F**) on splenic VPS34-deficient (Cre^+^) and VPS34-sufficient (Cre^-^) Treg cells from *Foxp3*^YFP-Cre/WT^*Pik3c3*^flox^ mosaic mice and *Foxp3*^YFP-Cre/WT^ control mice. **G – H**) mean fluorescence intensity (MFI) of CD80 on antigen-presenting cells (APCs) from *Foxp3*^YFP-Cre^*Pik3c3*^flox^ mice and *Foxp3*^YFP-Cre^ control mice (WT) (**G**) and *Foxp3*^YFP-Cre/WT^*Pik3c3*^flox^ mosaic mice and *Foxp3*^YFP-Cre/WT^ control mice (**H**). **I)** Representative flow cytometry plot for CD44 and CD62L expression and bar graph representing expression level of CD44 on splenic Treg cells from *Foxp3*^YFP-Cre^*Pik3c3*^flox^ mice and *Foxp3*^YFP-Cre^ control mice (WT). **J)** Expression level of CD44 on splenic VPS34-deficient (Cre^+^) and VPS34-sufficient (Cre^-^) Treg cells from *Foxp3*^YFP-Cre/WT^*Pik3c3*^flox^ mosaic mice. **K**) Representative flow cytometry plot for KLRG1 and CD44 expression and bar graph representing expression level of KLRG1on splenic Treg cells from *Foxp3*^YFP-Cre^*Pik3c3*^flox^ mice and *Foxp3*^YFP-Cre^ control mice (WT). **L**) Expression level of CD69, LAG-3, PD-1, and GITR on splenic VPS34-deficient (Cre^+^) and VPS34-sufficient (Cre^-^) Treg cells from *Foxp3*^YFP-Cre/WT^*Pik3c3*^flox^ mosaic mice. *Foxp3*^YFP-Cre^*Pik3c3*^flox^ mice and the respective control mice were between 4 and 5.5 weeks of age. *Foxp3*^YFP-Cre/WT^*Pik3c3*^flox^ mosaic mice and the respective control mice were between 8 and 12 weeks of age. n = 3-7 mice per group. Statistical significance was determined using an unpaired two-tailed Student’s t-test (**C, D, E, I**), paired two-tailed Student’s t-test (**A, B, F, G, H, J, K, L**). Results are pooled from 2 to 4 independent experiments.

Treg cells are characterised by the constitutive expression of the interleukin-2 (IL-2) receptor and CTLA-4. CTLA-4 on Treg cells captures CD80 and CD86 from APCs through a process termed transendocytosis, thus depriving Tcon of these costimulatory ligands (Omar S Qureshi et al., 2011). Similarly, the IL-2 receptor on Treg cells internalises and degrades IL-2, thus depriving Tcon of this essential cytokine (Pandiyan et al., 2007).

Expression of CTLA-4 (**Fig. 2C**) and the IL-2 receptor α-chain (also referred to as CD25; **Fig. 2D**) was increased on Treg cells from *Foxp3*^YFP-Cre^*Pik3c3*^flox^ mice. However, CTLA-4 and CD25 expression on VPS34-deficient (Cre^+^) Treg cells was indistinguishable from VPS34-sufficient Treg cells (Cre^-^) in *Foxp3*^YFP-Cre/WT^*Pik3c3*^flox^ mosaic mice (**Fig. 2E, F**, respectively). CTLA-4 expressed on Treg cells can reduce CD80 expression on APCs (Omar S Qureshi et al., 2011). Whereas CD80 was increased on splenocytes from *Foxp3*^YFP-Cre^*Pik3c3*^flox^ mice (**Fig. 2E**), CD80 levels on splenocytes from *Foxp3*^YFP-Cre/WT^*Pik3c3*^flox^ mosaic mice were not altered (**Fig. 2H**). To further investigate the phenotypical changes associated with VPS34-deletion in Treg cells, we analysed cell surface markers associated with Treg cell activation and function. VPS34-deficient Treg cells from *Foxp3*^YFP-Cre^*Pik3c3*^flox^ mice expressed elevated levels of the costimulatory receptor ICOS (**Fig. S2F**), while CD38 expression was decreased (**Fig. S2G**). However, we found no difference in the expression of these markers in VPS34-deficient (Cre^+^) compared to VPS34-sufficient (Cre^-^) Treg cells from *Foxp3*^YFP-Cre/WT^*Pik3c3*^flox^ mosaic mice (**Fig. S2H, I**, respectively). These results indicate that the differential expression of these markers is driven by the inflammatory environment. By contrast, the expression level of CD44 was both reduced in VPS34-deficient Treg cells from *Foxp3*^YFP-Cre^*Pik3c3*^flox^ mice (**Fig. 2I**) and among VPS34-deficient (YFP^+^ Cre^+^) Treg cells compared to VPS34-sufficient (YFP^-^ Cre^-^) Treg cells from *Foxp3*^YFP-Cre/WT^*Pik3c3*^flox^ mosaic mice (**Fig. 2J**). CD44 is a prominent marker for activation and reduced expression on VPS34-deficient Treg cells raised the hypothesis that loss of VPS34-kinase activity impairs the maturation of Treg cells into activated Treg cells or their survival. By contrast, expression of KLRG-1 was unchanged in VPS34-deficient (YFP^+^ Cre^+^) Treg cells compared to VPS34-sufficient (YFP^-^ Cre^-^) Treg cells from *Foxp3*^YFP-Cre/WT^*Pik3c3*^flox^ mosaic mice (**Fig. 2K**). In contrast to CD44, other activation markers such as CD69, LAG3, and PD-1 were significantly downregulated among VPS34-deficient (YFP^+^ Cre^+^) Treg cells compared to VPS34-sufficient (YFP^-^ Cre^-^) Treg cells from *Foxp3*^YFP-Cre/WT^*Pik3c3*^flox^ mosaic mice (**Fig. 2L**). CD69 is an early activation marker that modulates the balance between Th1/Th17 and Treg cells and has been proposed as a metabolic gatekeeper (Cibrián and Sánchez-Madrid, 2017). LAG3 is expressed after activation and is required for maximal regulatory activity of Treg cells (Zhang et al., 2017) and PD-1, through the PD-1/PD-L1 axis, regulates the differentiation and function of Treg cells (Francisco et al., 2009) . Interestingly, the expression of GITR, which is involved in Treg cells differentiation and expansion, was increased on VPS34-deficient Treg cells (**Fig. 2L**). It has been suggested that triggering of GITR decreases the suppressive activity of Treg cells (Ephrem et al., 2013).

The reduced proportion of Treg that expressed CD44, CD69, or PD1 data suggests VPS34 is intrinsically required for Treg cells to differentiate into a more mature phenotype and/or that mature Treg cells depend on VPS34 for their survival.

### VPS34-deletion does not interfere with known Treg cell suppressive functions

Next, we aimed to assess which suppression mechanisms were affected by loss of VPS34 in Treg cells. Given that VPS34 regulates several steps in the endolysosomal degradation process, we investigated whether loss of VPS34 kinase activity impairs endocytosis-mediated suppression mechanisms, i.e. the consumption of IL-2 or internalisation of the costimulatory ligands CD80/CD86.

First, we determined whether the IL-2 receptor signals normally on VPS34-deficient Treg cells. IL-2-stimulated STAT5 phosphorylation was unchanged between VPS34-deficient (YFP^+^ Cre^+^) and VPS34-sufficient (YFP^-^ Cre^-^) Treg cells from *Foxp3*^YFP-Cre/WT^*Pik3c3*^flox^ mosaic mice (**Fig. 3A**). To address the effect of VPS34-deficiency on the endocytosis of IL-2-IL-2 receptor complexes, we developed an assay to quantify the consumption of IL-2 by flow cytometry (**Fig. 3B**). In accordance with normal levels of CD25 on their surface (**Fig. 2G**), both cell surface-bound and internalised IL-2 levels did not differ between VPS34-deficient (YFP^+^ Cre^+^) and VPS34-sufficient (YFP^-^ Cre^-^) Treg cells from *Foxp3*^YFP-Cre/WT^*Pik3c3*^flox^ mosaic mice (**Fig. 3C**). Hence, both IL-2 receptor-mediated signalling and internalisation of IL-2 are unperturbed by VPS34 deficiency.

**Fig. 3.**
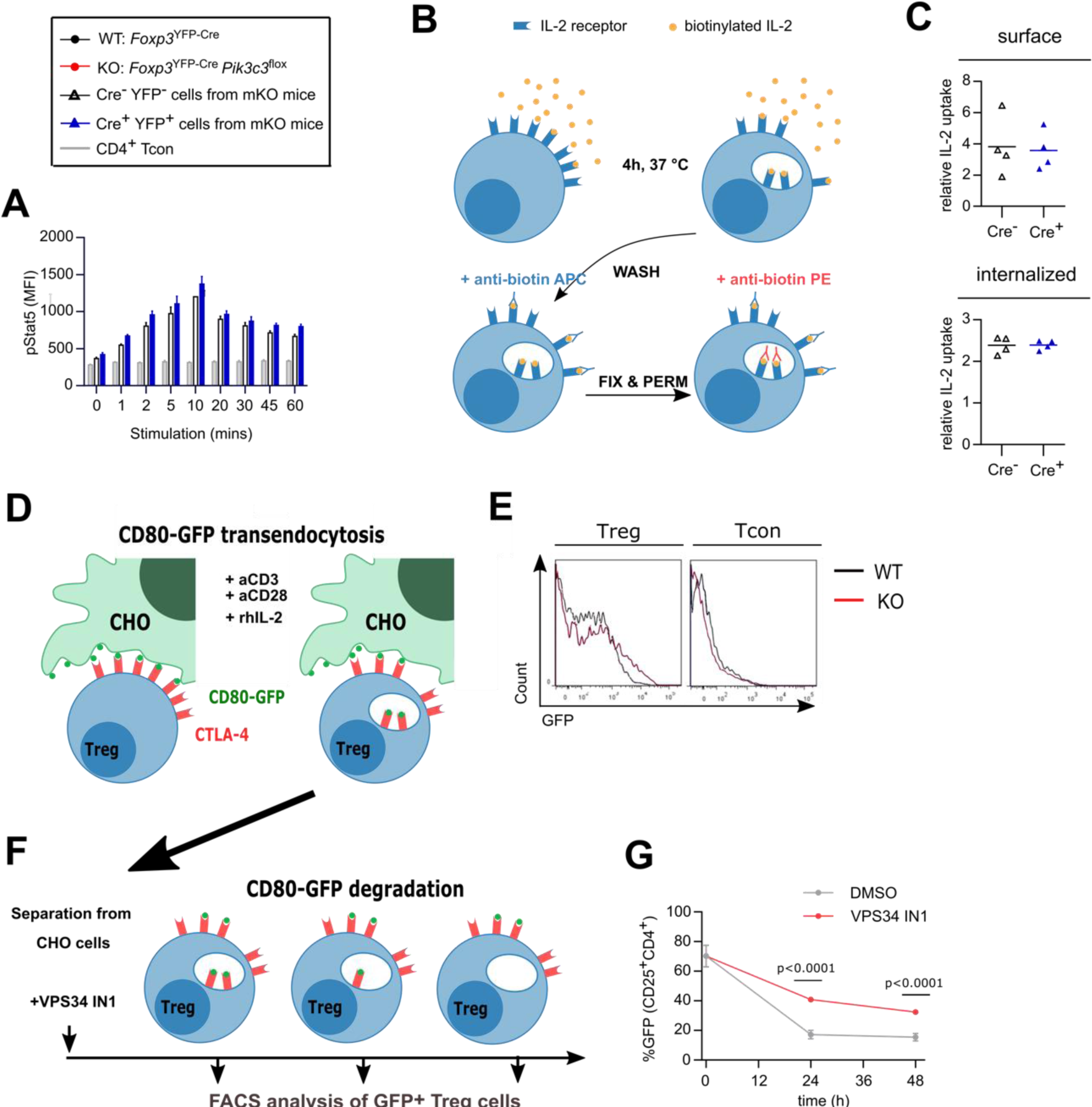
Treg cells from *Foxp3*^YFP-Cre^*Pik3c3*^flox^ mice efficiently internalise IL-2 but might displayed impaired degradation of CD80. **A)** Lymphocytes from *Foxp3*^YFP-Cre/WT^*Pik3c3*^flox^ mice were stimulated with 20 ng/ml rhIL-2 and stained for phospho-Stat5 for analysis by flow cytometry. The mean MFI of 3 mice is shown at each time point, and error bars indicate standard deviation. **B**) Schematic representation of the IL-2 internalisation assay. Lymphocytes were incubated at 37°C for 4 h in the presence of biotinylated rhIL-2. After washing, surface-bound and internalised IL-2 were identified by flow cytometry using anti-biotin antibodies in different conjugations. **C)** Median fluorescence intensity (MFI) of surface-bound and internalised IL-2 in Treg cells from mosaic *Foxp3*^YFP-Cre/WT^*Pik3c3*^flox^ and wild-type *Foxp3*^YFP-Cre/WT^ *Pik3c3*^WT^ mice. **D**) Schematic representation of the transendocytosis assay. Treg cells were enriched from *Foxp3*^YFP-Cre^*Pik3c3*^flox^ and *Foxp3*^YFP-Cre^ *Pik3c3*^WT^ mice and co-cultured for 24 h with an equivalent number of Chinese Hamster Ovary (CHO) cells expressing GFP-tagged CD80 on their surface. Tregs should internalise and accumulate CD80-GFP. **E**) Representative histograms showing GFP-tagged CD80 fluorescence associated on Treg cells from *Foxp3*^YFP-Cre^*Pik3c3*^flox^ and *Foxp3*^YFP-Cre^ *Pik3c3*^WT^ mice. **F**) Schematic representation of the pulse-chase assay. Similar to in (**D**), Treg cells from *Foxp3*^YFP-Cre^ *Pik3c3*^WT^ mice were enriched and co-cultured for 24 h with a 10:1 ratio of Chinese Hamster Ovary cells expressing GFP-tagged CD80 on their surface. After 24h co-culture, Treg cells were separated from the rest of the cells by fluorescence-activated cell sorting (FACS) and cultured presence of VPS34 IN1, a selective Vps34 inhibitor, for up to 48 h. **G**) The percentage of the GFP signal was assessed by flow cytometry directly after separation from CHO cells (0 h), and then 24 and 48 h after separation. *Foxp3*^YFP-Cre^*Pik3c3*^flox^ mice and wild-type control mice were between 4 and 5.5 weeks of age. *Foxp3*^YFP-Cre/WT^*Pik3c3*^flox^ mosaic mice and wild-type control mice were between 8 and 12 weeks of age. n = 3-4 mice per group. Statistical significance was determined using an unpaired two-tailed Student’s t-test. Statistical significance between YFP-Cre^-^ and YFP-Cre^+^ Treg was determined for each time point using an unpaired t-test; no significant difference was found at any point. Results are pooled from two to three independent experiments.

We next assessed the ability of VPS34-deficient Treg cells to acquire CD80 from APCs, using a previously described assay (Omar S. Qureshi et al., 2011) (**Fig. 3D**). VPS34-deficient Treg cells were found to acquire more CD80-GFP than WT Treg cells, most likely reflecting elevated CTLA-4 expression (**Fig. 2C**), while Tcon did not appear to acquire GFP (**Fig. 3E**).

Following transendocytosis, the internalised cargo is degraded through the endolysosomal pathway, a system relying on PI3P for the recruitment of endosomal sorting complexes (Bilanges et al., 2019). Therefore, we considered whether the delivery of endosomal cargo to lysosomes for degradation is impaired when VPS34 is inactivated, despite intact initial internalisation. Indeed, we found that VPS34-IN1, a selective small molecule inhibitor of VPS34 (Bago et al., 2014), delayed the degradation of the internalised CD80 cargo as evidenced by higher GFP fluorescence in Treg cells after separation from the donor cells (**Fig. 3F, G**). Together, these results suggest that VPS34 inhibition, resulting in reduced PI3P levels, impairs the degradation of the internalised ligand in the lysosomes.

### Defective autophagy may contribute to, but is not sufficient to explain the profound autoimmune disease in *Foxp3^YFP^*^-Cre^*Pik3c3^flox^*mice

Autophagy, a highly conserved degradation process critical to maintain cellular homeostasis, plays an important role in establishing Treg cell-mediated immune tolerance, supporting lineage stability and survival fitness (Wei et al., 2016). During the formation of the autophagic membrane, the protein LC3 (also known as ATG8) becomes lipidated and is referred to as LC3-II (to differentiate it from the non-lipidated form LC3-I). Autophagy is increased after the initial activation of T cells, potentially as a mechanism to degrade the increasing cytoplasmic content that is generated as T cells become metabolically more active (Dowling and Macian, 2018). Rather than the expected decrease in LC3-II abundance in VPS34-deficient Treg cells, we observed a mild increase (**Fig**. **4A**). This suggests that VPS34 deficiency in Treg cells does not affect autophagy initiation, though we cannot exclude an impact of VPS34-deficiency on autophagy *in vivo* under conditions of stress or nutrient deprivation.

**Fig. 4.**
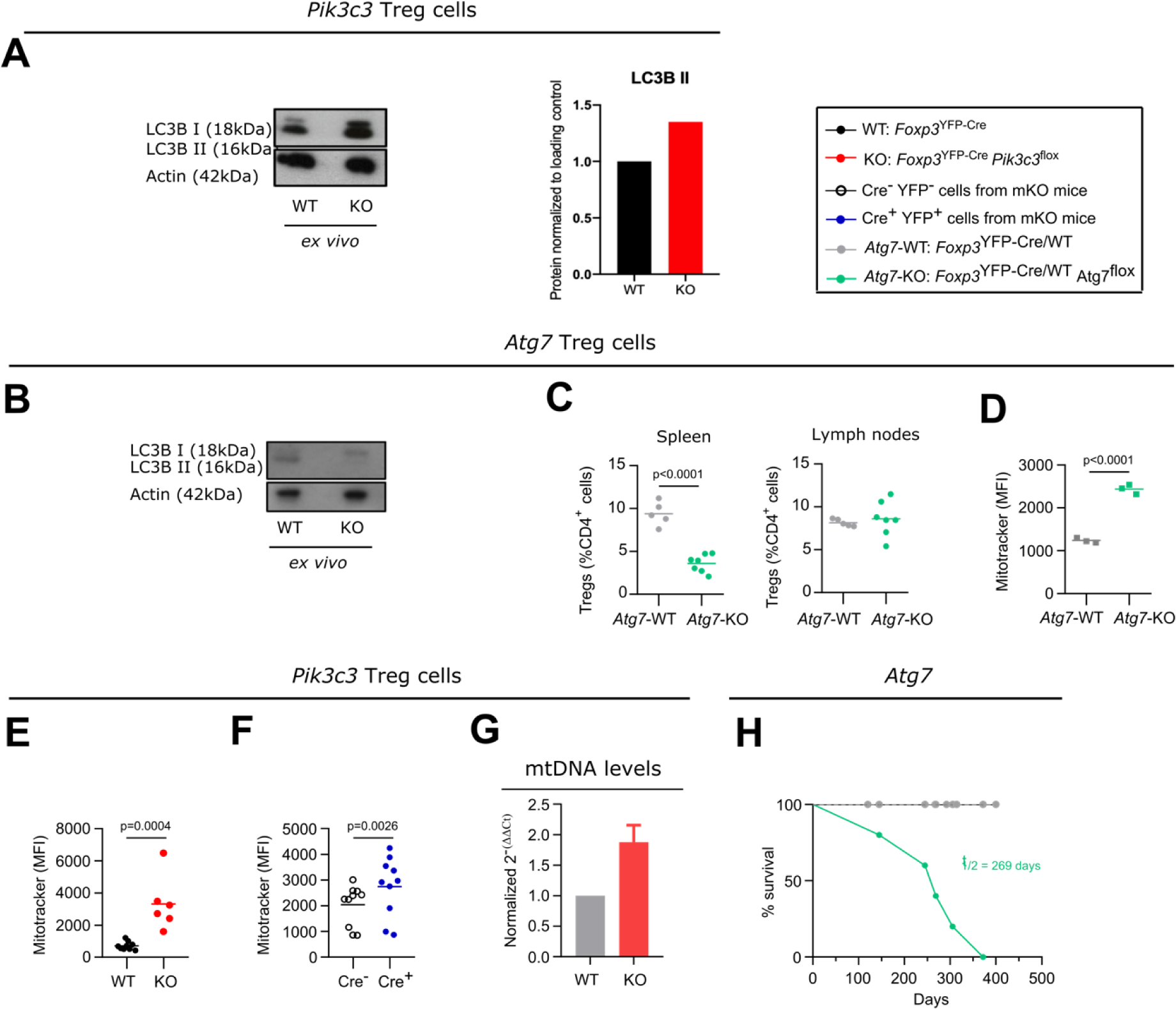
Defective autophagy in VPS34-deficient Treg cells might contribute to, but is not sufficient to explain the phenotype of *Foxp3*^YFP-Cre^*Pik3c3*^flox^ mice. **A – B**) Densiometry of LC-3 I and LC-3II in *ex vivo* Treg cells from *Foxp3*^YFP-Cre^*Pik3c3*^flox^ (**A**) or *Foxp3*^YFP-Cre^*Atg7*^flox^ mice (**B**) and their respective controls (WT) was determined by Western blot analysis. Actin was used as a loading control. Amounts of all proteins were normalized to actin and are relative to amounts in WT cells. **C**) Percentage of Treg cells from the spleen and the lymph nodes of *Foxp3*^YFP-Cre^ *Atg7*^flox^ mice and wild-type control mice (WT). **D – F**) Mean fluorescence intensity (MFI) of the MitoTracker Orange dye in splenic Treg cells from *Foxp3*^YFP-Cre^ *Atg7*^flox^ (**D**), *Foxp3*^YFP-Cre^*Pik3c3*^flox^ (**E**), mosaic *Foxp3*^YFP-Cre/WT^*Pik3c3*^flox^ (**F**) mice, and the respective wild-type mice (WT). **G)** Quantitative PCR analysis of the mitonchondria DNA (mtDNA) levels of FACS-sorted Treg cells from *Foxp3*^YFP-Cre^*Pik3c3*^flox^ and control mice (WT). **H**) Survival curves for *Foxp3*^YFP-Cre^*Atg7*^flox^ (green) and wild-type *Foxp3*^YFP-Cre^ *Atg7^WT^* mice (grey). *Foxp3*^YFP-Cre^*Pik3c3*^flox^ mice and wild-type control mice were between 4 and 5.5 weeks of age. *Foxp3*^YFP-Cre/WT^*Pik3c3*^flox^ mosaic mice, *Foxp3*^YFP-Cre^*Atg7*^flox^, and the respective control mice were between 8 and 12 weeks of age. n = 3-11 mice per group. Statistical significance was determined using a two-tailed Student’s t-test. Results are pooled from 2 to 3 independent experiments.

In order to more directly assess the role of autophagy in Treg cells *in vivo*, we deleted ATG7, a protein essential for the induction of autophagy (Komatsu et al., 2005), specifically in Treg cells by crossing *Foxp3*^YFP-Cre^ mice to an *Atg7*^flox^ strain. This lack of ATG7 in Treg cells led to impaired LC3-II lipidation, consistent with compromised autophagy initiation (**Fig. 4B**), as well as a reduced percentage of Treg cells in the spleen but not the lymph nodes (**Fig. 4C**). This observation suggests that more mature Treg cells, as predominantly present in the spleen compared to the lymph nodes, rely more on autophagy compared to more naïve Treg cells.

Cellular metabolism is important for Treg cell homeostasis and function. As part of the maintenance of cellular homeostasis, mitophagy, a selective form of autophagy required for the removal of defective and excessive mitochondria, plays a critical role. Autophagy-deficient cells accumulate damaged mitochondria with altered membrane potential (Pua et al., 2009). Accordingly, we detected more mitochondria in ATG7-deficient Treg cells (**Fig. 4D**). Similarly, we observed an increase in the staining for mitochondria in VPS34-deficient Treg cells from *Foxp3*^YFP-Cre^*Pik3c3*^flox^ mice and YFP^+^ Cre^+^ Treg cells from *Foxp3*^YFP-Cre/WT^*Pik3c3*^flox^ mosaic mice (**Fig. 4E, F**, respectively). Since the incorporation of the used dye is not only dependent on mitochondrial potential, but also on mitochondrial mass, we assessed the level of mitochondrial DNA in Treg cells from *Foxp3*^YFP-Cre^*Pik3c3*^flox^ mice by quantitative PCR analysis. Results showed elevated mitochondrial DNA (**Fig. 4G**), suggesting an increase in mitochondrial number and impaired intracellular mitochondrial clearance, consistent with a defect in autophagy in VPS34-deficient Treg cells. However, in contrast to *Foxp3*^YFP-Cre^*Pik3c3*^flox^ mice, *Foxp3*^YFP-Cre^*Atg7*^flox^ mice did not show a Scurfy-like phenotype and survived for up to a year (**Fig. 4H**, t½ = 269 days), consistent with results published by Wei *et al*. (Wei et al., 2016). We therefore concluded that defective autophagy may contribute to, but is not sufficient to explain, the profound autoimmune lymphoproliferative disease observed in *Foxp3*^YFP-Cre^*Pik3c3*^flox^ mice. Moreover, the presence of LC3-II in VPS34-deficient, but not ATG7-deficient, Treg cells suggests that VPS34 deletion is not essential for autophagy induction, but may control selective forms of autophagy such as mitophagy.

### Loss of VPS34 kinase activity increases cellular respiration in Treg cells

VPS34 is primarily thought to affect trafficking of subcellular vesicles and their associated proteins. A block at any stage in these processes could alter the steady-state levels of proteins involved in steps before and after the block. Therefore, with the aim to discover such novel suppressive functions of Treg cells, we performed quantitative high-resolution mass spectroscopy to compare the proteomic profile of VPS34-deficient and VPS34-sufficient Treg cells (Fig. 5A). To exclude potential secondary effects caused by systemic inflammation, we performed the analysis on YFP+ Cre+ VPS34-deficient Treg cells from phenotypically normal Foxp3YFP-Cre/WTPik3c3flox mosaic mice and used YFP+ Cre+ VPS34-sufficient Treg cells from Foxp3YFP-Cre/WTPik3c3WT mice as a control. We detected about 6,000 distinct proteins, demonstrating that we can effectively isolate sufficient numbers of primary Treg cells from mice for comprehensive proteomic analyses. We performed statistical analyses to assess differences in total protein content and identified a total of 157 differentially-expressed proteins, 124 of which were more abundant and 33 less abundant in VPS34-deficient Treg cells compared to VPS34-sufficient Treg cells (Fig. 5B).

**Fig. 5.**
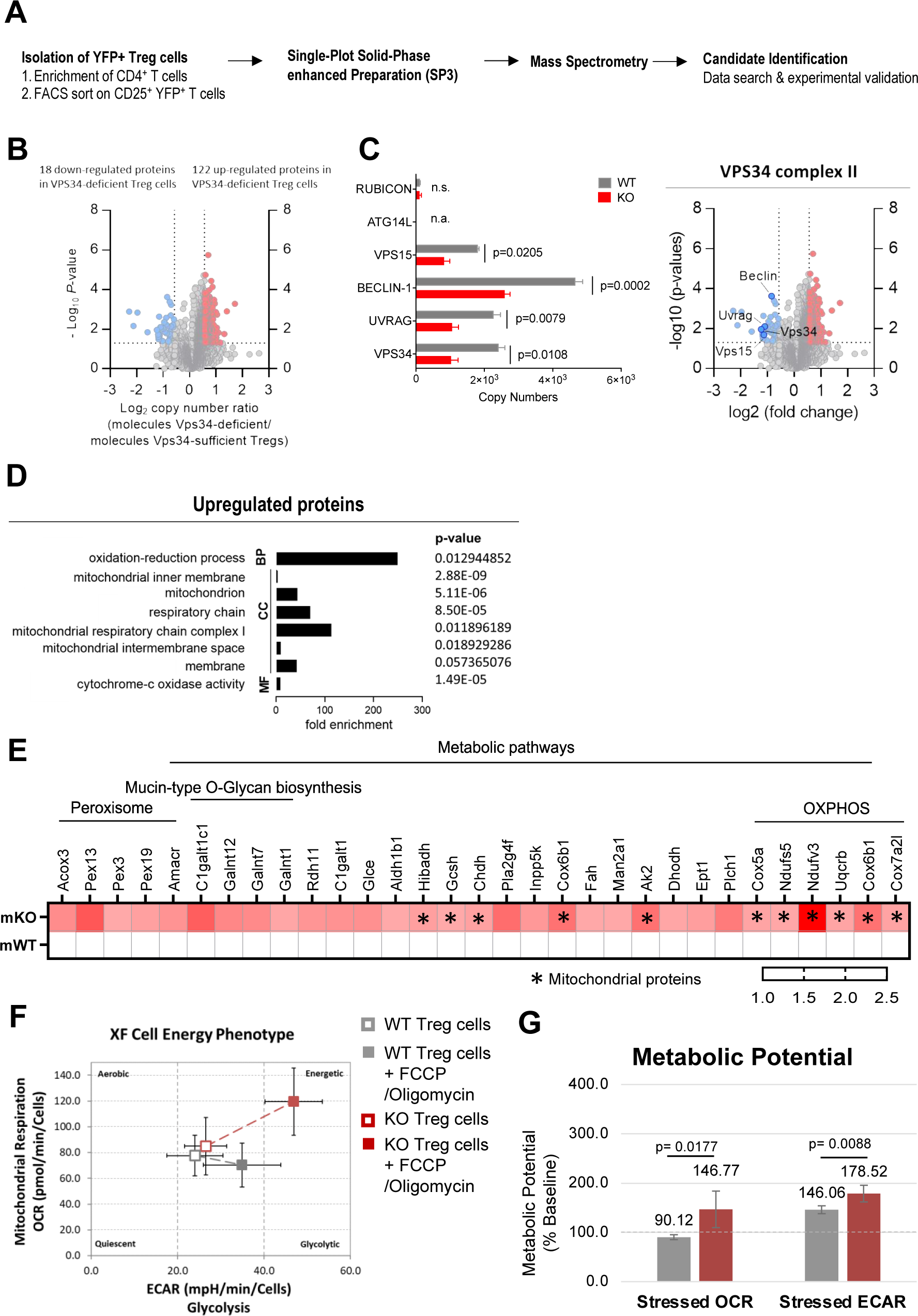
Proteomic profiling revealed that loss of VPS34 kinase activity increases cellular respiration in Treg cells. **A**) Workflow for the proteomic profiling of Treg cells from *Foxp3*^YFP-Cre/WT^*Pik3c3*^flox^ mosaic mice and *xp3*^YFP-Cre/WT^ *Pik3c3*^WT^ control mice. **B**) Volcano plot representing the ratio of the protein copy numbers in VPS34-deficient Treg cells compared to VPS34-sufficient Treg cells. Proteins that exhibited statistically significant increase in abundance (p < 0.05 by one-sample Student’s t test) by greater than 1.5-fold are shown in red, while proteins that exhibit statistically significant reduction in abundance are shown in blue. **C**) Volcano plot highlighting the binding partners of the VPS34 complex II. Bar graphs showing estimated copy numbers calculated using the proteomic data for RUBICON, ATG14L, VPS15, BECLIN-1, UVRAG, and VPS34. Individual data points from the three biological replicates performed for the proteomic analysis are shown, with the bar showing the means ± S.D. *p* values were calculated by two-tailed, one-sample Student’s *t* test. **D**) Analysis of biological processes (BP), cellular components (CC), molecular function (MF) involving proteins upregulated in VPS34-deficient Treg cells identified in the proteomic profiling analysis. **E**) Heat map presenting proteins upregulated in VPS34-deficient Treg cells identified in the proteomic profiling analysis. **F**) Oxygen consumption rate (OCR) and extracellular acidification rate (ECAR) of VPS34-deficient Treg cells before and after addition of 0.5 µM FCCP and 10μM oligomycin following the Seahorse XF Cell Energy Phenotype Test Assay protocol. **G**) Percentage increase of stressed OCR over baseline OCR, and stressed ECAR over baseline ECAR. The metabolic potential is the measure of cells’ ability to meet an energy demand via respiration and glycolysis. *Foxp3*^YFP-Cre/WT^*Pik3c3*^flox^ mosaic mice and wild-type control mice were between 8 and 12 weeks of age. Statistical significance was determined using an unpaired two-tailed Student’s t-test. Results are pooled from 2 to 3 independent experiments.

The expression of VPS34 and its associated proteins VPS15, BECLIN-1, and UVRAG (specific to VPS34-complex 2) were reduced by half (Fig. 5C), possibly resulting from the reduced expression of the truncated VPS34Δ21 protein, affecting the VPS34 complex integrity. Interestingly, we did not detect the VPS34 complex 1 protein ATG14L, which may be found at low levels in resting Treg cells. Levels of Run domain Beclin-1 interacting and cysteine-rich containing protein (RUBICON) which replaces ATG14L during non-canonical autophagy 29, were also low and unaffected by VPS34-deficiency.

Using DAVID functional annotation clustering to identify biologically relevant groups of genes with altered expression, we found that the loss of VPS34-kinase activity led to downregulation of proteins involved in multiple distinct molecular functions and biological processes. Meanwhile, the majority of the upregulated proteins were involved in biological processes relating to oxidation-reduction processes and energy metabolism (Fig. 5D). Up-regulated proteins related to cellular components involved mitochondrial functions, specifically mitochondrial inner membrane, mitochondrion, respiratory chain, mitochondrial respiratory chain complex I, mitochondrial intermembrane space, and membrane proteins, as well as cytochrome c-oxidase activity (Fig. 5D). These data suggested that VPS34-deficiency results in an increased abundance of proteins involved in mitochondrial function and energy metabolism, correlating with the elevated mitochondria mass observed in VPS34-deficient Treg cells (Fig. 4E, F). Specifically, we identified increased levels of proteins involved in metabolic pathways (Fig. 5E), and OXPHOS (Fig. 5F) with subunits of the electron transport chain complexes I, III and IV, namely Uqcrb, Cox6b1, Cox7a2l, Cox5a, Ndufs5, and Ndufv3 being the most significantly affected. The increased abundance of proteins involved with OXPHOS may in part reflect the increased number of mitochondria found in VPS34-deficient Treg cells (Fig 4G).

These findings led to the hypothesis that the loss of VPS34 kinase activity induces a state of metabolic dysregulation in Treg cells. We therefore assessed whether differences in protein levels identified by proteomic profiling in VPS34-deficient Treg cells were accompanied by biologically relevant differences in energy levels. To quantify mitochondrial respiration in VPS34-deficient Treg cells, we measured the rate of mitochondrial respiration in the cells (by measuring the oxygen consumption rate (OCR), the rate of decrease of oxygen concentration in the assay medium) and the rate of glycolysis of the cells (by measuring the extracellular acidification rate (ECAR), the rate of increase in proton concentration (or decrease in pH) in the assay medium). This assay allows simultaneous measurement of the two major energy-producing pathways of the cell - mitochondrial respiration and glycolysis - by the simultaneous addition of oligomycin and FCCP. Oligomycin inhibits the production of ATP by mitochondria, and causes a compensatory increase in the rate of glycolysis as the cells attempt to meet their energy demands via the glycolytic pathway. FCCP depolarizes the mitochondrial membrane and increases oxygen consumption as the mitochondria attempt to restore the mitochondrial membrane potential. We found that VPS34-deficient Treg cells displayed a slight increase in both OCR and ECAR (Fig. 5F) and an increased metabolic potential (Fig. 5G). These data indicate that VPS34 may restrict the glycolytic and respiratory potential of Treg, possibly by increasing mitophagy. Given that Treg cells tend to be more dependant on fatty acid oxidation and are less metabolically active than Teff cells, this increased metabolic activity may paradoxically also limit their suppressive phenotype(capacity).

## Discussion

The class III PI3K VPS34 plays a fundamental role in endocytosis, intracellular vesicular trafficking, and autophagy – key processes that control T cell function. Autophagy is critical for regulating T cell activation and differentiation by degrading cytoplasmic components, where the degraded material is recycled and used by cellular metabolic pathways, linking autophagy to metabolism (Kabat et al., 2016; Wei et al., 2016).

We found that mice with VPS34 inactivated in Treg cells died within 6 weeks of birth from a lymphoproliferative disease, similar to what has previously been observed in *Foxp3* knockout mice displaying the *Scurfy* phenotype (Brunkow et al., 2001). However, in contrast to *Foxp3* knockout mice, and most other models that recapitulate the *Scurfy* phenotype, VPS34-deficient Treg cells developed and populated the peripheral lymphoid organs, suggesting that loss of VPS34 affects Treg cell suppressive functions rather than survival. However, neither the suppression of Tcon nor the consumption of IL-2 and subsequent phosphorylation of STAT-5, were affected by the lack of VPS34. VPS34-deficient Treg cells were also capable of removing CD80 from target cells by the process of transendocytosis. PI3P is present within the endolysosomal system and particularly predominant in early endosomes where it is required for the recruitment of PX- and FVYE-domain containing proteins found in several endosomal sorting machineries, such as retromer and ESCRT (Backer, 2016). Inhibition of VPS34, and subsequent loss of PI3P, prevents the recruitment of Armus, a Rab7 GAP, resulting in Rab7 hyperactivation, enlarged endosomes and altered endosomal dynamics (Jaber et al., 2016). Indeed, the degradation of the endocytosed CD80 cargo was impaired by inhibition of VPS34. This may have a greater impact on Treg cells *in vivo* as their overall processive capacity to endocytose CD80 would be impaired.

VPS34 is also essential for autophagy. However, conditional deletion of *Atg7* or *Atg16l*, both of which are essential for autophagy, in Treg cells led to a less severe phenotype than deletion of VPS34 (**Fig. 4H** and (Kabat et al., 2016; Wei et al., 2016)). Moreover, we did not observe severe impact on LC3-II formation in VPS34-deficient Treg cells. However, we cannot exclude the possibility that particular stimuli or environmental cues may induce autophagy in a VPS34-dependent manner. Therefore, impairment of autophagy might contribute by perturbing the homeostasis of VPS34-deficient Treg cells; however, such loss of autophagy is not sufficient to explain the phenotype observed in mice with VPS34-deficient Treg cells.

The inflammatory environment present in *Foxp3*^YFP-Cre^*Pik3c3*^flox^ mice might impact the phenotype of Treg cells in a manner that is not a direct consequence of intrinsiv VPS34-defciciency. We therefore generated mice with a ‘mosaic’ deletion of VPS34 in Treg cells. Indeed, these mice were healthy and did not show any sign of an inflammatory disease, demonstrating that VPS34-deficient Treg cells do not actively cause pathology, or, if they did, that the VPS34-sufficient Treg cells can effectively suppress them. The proportion of Treg cells expressing CD44 was lower among VPS34-deficient Treg cells from mosaic mice, similar to what we have also observed when VPS34 was knocked-out in all Treg cells, suggesting that VPS34 is intrinsically required for Treg cells to differentiate into a more mature phenotype or, alternatively, that mature Treg cells depend on VPS34 for their survival.

In an attempt to identify and characterise Treg cell suppression mechanisms that are affected by the loss of VPS34, we performed proteomic profiling. A large proportion of aberrantly expressed proteins were associated with OXPHOS. This could at least in part be explained by increased abundance of mitochondria as a consequence of reduced mitophagy. This increase in cellular metabolism was associated with increased mitochondrial respiratory complexes. A number of recent reports have shown that the numbers of Treg cells and their phenotypical plasticity are regulated by metabolic processes ^31^. Treg cells are known to have a ‘‘metabolic edge’’ for survival through their bias toward mitochondrial respiration, and mitochondrial metabolism supports both their immunosuppressive functions as well as survival in lactate-rich environments ^32^. Indeed, a recent report correlated increase in mitochondrial oxidative stress in Treg cells being an underpinning cause of autoimmunity ^33^. Others have investigated the relationship between Treg cell fitness, suppressive functions, and metabolic activity show that mice developed a fatal inflammatory disorder similar to the one observed in *Foxp3*^Cre-YFP^*Pik3c3*^flox^ mice. However, this phenotype was associated with impaired mitochondrial function and cellular metabolism, rather than increased metabolic activity as we find in VPS34-deficient Treg cells ^34–37^. Nevertheless, together these data highlight that altered mitochondrial metabolism can profoundly affect Treg function.

Further evidence suggests that FOXP3 itself directs reprograming of T cell metabolism by suppressing *Myc* and glycolysis, while enhancing OXPHOS, suggesting FOXP3-targets may contribute to the regulation of mitochondrial respiratory complexes to ensure preservation of Treg cell function ^32^. In *Foxp3*^Cre-YFP^*Pik3c3*^flox^ mice and *Foxp3*^Cre-YFP/WT^*Pik3c3*^flox^ mosaic mice, expression of *Foxp3* was unchanged (*data not shown*), suggesting that VPS34 influences metabolic reprogramming by limiting the spare respiratory cacacity, possibly by promoting mitophagy. How this affects the fitness VPS34-deficient function remains unresolved.

This work reveals a fundamental and novel role for the class III PI3K VPS34 in the maintenance of Treg cell suppressive mechanisms, and has drawn a link between VPS34 and Treg cell metabolism. While defects in endocytosis or autophagy alone may not be sufficient to explain the profound phenotype of mice with VPS34-deficient Treg cells, a more general failure to process endocytic and/or autophagic intracellular vesicles and their cargo may combine to render VPS34-deficient Treg cells incapable of suppressing autoimmune responses *in vivo*.

## Materials and Methods

### Mice

All procedures involving mice were carried out in accordance with the United Kingdom Home Office regulations (Animals (Scientific procedures) Act 1986, PPLs 70-7661, P0AB4361E) and reviewed by the local Animal Welfare and Ethics Review Bodies. Mice were maintained in individually-ventilated cages under specific pathogen-free conditions at the Babraham Institute’s Biological Services Unit, and the Biological Service Units of the University of Cambridge. We used male and female animals aged between 3.5 to 52 weeks in all experiments.

*Pik3c3*^flox^ mice were generated by flanking exon 21 of the *Pik3c3* gene with *loxP* recombination sites. *Foxp3*^YFP-Cre^ mice expressing a YFP-Cre protein knocked into the 3’ untranslated region of the *Foxp3* gene (Rubtsov et al., 2008) were a kind gift from Dr. Alexander Rudensky. Both strains were crossed to obtain mice with cell type-specific deletion of VPS34 (*Foxp*^YFP-Cre^*Pik3c3*^flox^ mice); *Foxp3*^YFP-Cre^ mice were used as wild-type controls. The location of *Foxp3* on the X chromosome enabled the breeding of heterozygous *Foxp3*^YFP-Cre/WT^ female offspring with both VPS34-deficient and VPS34-sufficient Treg cells (mosaic *Foxp3*^YFP-Cre/WT^*Pik3c3*^flox^ mice). Treg cell-specific deletion of exon 21 of *Pik3c3* was confirmed by PCR analysis from sorted YFP^+^ Treg cell populations from *Foxp3*^YFP-Cre^*Pik3c3*^flox^ mice and *Foxp3*^YFP-Cre^ control mice (Supp Fig. 1D). *Foxp3*^YFP-Cre^ mice were also crossed to *Atg7*^flox^ mice to obtain autophagy-deficient Treg cells.

### Buffers and Media

**Cell staining buffer (CSB):** PBS/2% foetal calf serum (FCS)

**T cell medium (TCM)**: RPMI-1640 (Gibco) with 5% v/v FCS, 0.005% v/v β-mercaptoethanol (Sigma), 1% v/v Pen Strep (Gibco), and 2mM L-glutamine (Gibco).

**T cell isolation buffer:** PBS/2% FCS.

**Cell Lysis Buffer:** 50 mM HEPES, 150 mM NaCl, 10 mM NaF, 10 mM Indoacetamide, 1% IGEPAL complemented with 1x of reconstituted 10x Complete Mini Protease Inhibitor Cocktail Tablet (Roche).

### Preparation of single cell suspensions from mouse tissues

Mice were culled by CO_2_ inhalation, followed by cervical dislocation; dissected tissues were kept on ice while processed. Single-cell suspensions were prepared from spleens, thymi and lymph nodes by pushing the tissue through 40 µm cell strainers (BD) using a syringe plunger. The cell suspensions were washed once with 5 ml cold PBS, and red blood cells in spleen samples were lysed using hypotonic ammonium chloride RBC lysis buffer (Sigma) according to the manufacturer’s instructions. Cells were washed once in 5ml cold PBS and collected by centrifugation. Cells were then resuspended at 1-3x10^6^ cells per sample and stained for flow cytometry as described below.

### T cell isolation

Negative selection of CD4^+^ T cells from peripheral lymph node cell suspensions were performed by using either the magnetic-activated cell sorting (MACS) Miltenyi kit or the mouse CD4 EasySep® (StemCell) kit. All incubation and wash steps were performed in T cell isolation buffer unless otherwise indicated. When using the Miltenyi kit, cells were resuspended at 1x10^8^ cells/ml and FITC-conjugated antibodies against mouse MHC-II, CD25, B220, CD8a, CD49b, and CD11b were added at a final dilution of 1:500 followed by incubation for 30 min at 4°C. Negative selection was performed using anti-FITC Microbeads (Miltenyi) and LS magnetic columns (Miltenyi) according to the manufacturer’s instructions, and unlabelled cells in the flow-through were collected. When using the StemCell kit, cells were isolated according to the manufacturer’s instructions. Regular purity checks by flow cytometry confirmed enrichment to >95% CD4^+^ T cells.

### T cell suppression assay

YFP^+^ Treg cells were isolated by sorting lymph node samples pre-enriched for CD25^+^ cells using a mouse CD25 Microbead kit (Miltenyi Biotec) on a FACSAria (BD Biosciences). CD4^+^CD25^-^ Tcon were isolated from the CD25^-^ fraction obtained from the same samples by negative selection using FITC-conjugated antibodies and anti-FITC Microbeads (Miltenyi Biotec) as described above (T cell isolation). Treg cells were co-cultured in known ratios with 1x10^5^ Tcon cells per well in 96-well round-bottom Nunclon plates (Nunc) and stimulated with 2x10^4^ anti-CD3/anti-CD28-coated Dynabeads (Dynal). After 96 h incubation at 37°C in an atmosphere of 5% CO_2_, proliferation was measured by [^3^H]-thymidine incorporation. [^3^H]-thymidine was added at 0.5μl (∼0.5 μCi) per well and plates incubated for 6 h before cells were harvested to UniFilter plates (Perkin Elmer) using a Tomtec 96 Harvester. Collection plates were air-dried overnight and 30 μl MicroScint™-20 (Perkin Elmer) added to each well. A TopcountTM scintillation counter (Perkin Elmer) was used to read the relative incorporation of [^3^H]-thymidine in each well.

### Enzyme-linked immunosorbent assay (ELISA)

ELISAs for interferon-γ (IFNγ) and interleukin-2 (IL-2) were performed using the Mouse IFNγ ‘Femto-HS’ Ready-SET-Go!^TM^ and Mouse IL-2 Ready-SET-Go!^TM^ ELISA kits (eBioscience) in accordance with the manufacturer’s instructions. Briefly, MaxiSorp plates (Nunc) were coated overnight at 4°C with capture antibody. All subsequent incubations occurred at room temperature and in the dark. Plates were washed in ELISA wash buffer (0.1% Tween 20 in PBS), and blocked with assay diluent for 1 h. Assay diluent was removed, and samples and standard curves were added to plates in duplicate or triplicate. Plates were incubated for 2 h, washed, and the detection antibody was added to all wells. After incubation for 1 h, plates were washed and incubated with avidin-HRP for 30 min. Plates were washed thoroughly, and tetramethylbenzidine substrate (TMB) added for 5-15 min until a discernible colour change had taken place. Reactions were stopped with 2N H_2_SO_4_ (Sigma), and plates read at 450 nm minus 570 nm on a Model 680 Microplate Reader (BioRad).

### IL-2 internalisation assay

Treg cells at a density of 10^5^ cells/well were resuspended in 25 μl TCM, and 10 μl biotinylated rhIL-2 or biotinylated control protein (from Fluorokine® Biotinylated Human IL-2 Kit, R&D Systems) added to each well. Plates were incubated at 37°C, 5% CO_2_ for 4 h, and cells washed once in cell staining buffer. Cells were stained for flow cytometry, initially for surface antigens including bound rhIL-2, using an APC-conjugated anti-biotin antibody (Miltenyi Biotech). Intracellular staining was then performed using a PE-conjugated anti-biotin antibody (Miltenyi Biotech) to detect internalised rhIL-2. Samples were analysed by flow cytometry.

### CD80 transendocytosis and pulse-chase assay

To test the capacity to deplete and digest CD80 from antigen-presenting cells (APCs) through CTLA-4–mediated transendocytosis, Treg cells were co-cultured with Chinese Hamster Ovary (CHO) cells expressing GFP-tagged CD80 (Omar S Qureshi et al., 2011) in CHO cell medium (DMEM containing 25 mM D-glucose (Sigma), 1 mM sodium pyruvate (PAA), 10% v/v FBS, 2 mM glutamine, 1% v/v Pen/Strep, 5 mM β-mercaptoethanol and 500 µg/ml G418 sulphate (Invitrogen)). CHO cells were co-cultured in a 1:10 ratio with CD4^+^ T cells (obtained by magnetic bead enrichment) in αCD3-coated (1 µg/ml) plates in CHO cell medium supplemented with rhIL-2 (20 ng/ml) and αCD28 (1 µg/ml). After 24 h co-culture at 37°C, 5% CO_2_, Treg cells were separated from the CHO cells by FACS sorting, transferred to a new plate and treated with the selective inhibitor VPS34-IN1 (1µM). The percentage and the MFI of GFP^+^ Treg cells was assessed by flow cytometry at the indicated time points.

### Flow cytometry

Single cell suspensions (1-3x10^6^ cells per sample) were stained for flow cytometry. For detecting cell surface antigens, cells were washed once and incubated for 30 min at 4°C with an antibody master mix prepared in cell staining buffer. Cells were washed once and fixed using a 4% paraformaldehyde solution. When biotinylated antibodies were used, cells were stained for 15 min at 4°C with fluorophore-conjugated streptavidin prior to fixation. For the detection of intracellular cytokines, cells were resuspended in T cell culture medium, and stimulated with 50ng/ml PdBU (Tocris), 1µM ionomycin (Sigma) and Brefeldin A (eBioscience) for 4 to 6 h at 37°C. Intracellular staining for the detection of *Foxp3* was performed using the eBioscience Foxp3/Transcription Factor Staining Buffer Set, according to the manufacturer’s instructions. Intracellular staining for the detection of cytokines was performed using the Biolegend Intracellular Staining Permeabilisation/Wash buffer, according to the manufacturer’s instructions. The Annexin V/Dead Cell Apoptosis Kit (BioLegend) was used according to manufacturer’s instructions to detect apoptotic cells.

To stain for mitochondrial potential/active OXPHOS, cells were incubated with 0.25 µM MitoTracker Orange CM-H_2_-TMRos or MitoTracker Orange CM-TMRos in RPMI-1640 medium containing 10% FCS and 1% Pen/Strep for 15 min at 37 ⁰C. Samples were analysed using BD LSRIIFortessa/Fortessa5, Cytek Aurora or Life Technologies Attune flow cytometers; data were analysed in FlowJo, Tree Star. Flow cytometry antibodies are summarised in Table I.

**Table 1:** Flow cytometry antibodies.

| Specificity | Clone | Conjugation | Concentration (μg/ml) | Supplier |
| --- | --- | --- | --- | --- |
| CD4 | RM4-5 | BV510 | 0.5 | BioLegend |
| CD4 | GK1.5 | BUV805 | 0.5 | BioLegend |
| CD8α | 53.6.7 | BV785 | 0.5 | BioLegend |
| CD8α | 53.6.7 | eFluor450 | 1 | eBioscience |
| CD8α | 53.6.7 | PerCP-Cy5.5 | 1 | eBioscience |
| CD11c | N418 | Pacific Blue | 1 | BioLegend |
| CD11c | N418 | APC-Cy7 | 1 | BioLegend |
| CD25 | PC61 | BV605 | 0.5 | BioLegend |
| CD25 | PC61 | BV785 | 0.5 | BioLegend |
| CD38 | 90 | PE Dazzle594 | 1 | BioLegend |
| CD38 | 90 | APC | 0.5 | BioLegend |
| CD44 | IM7 | PerCP Cy5.5 | 0.5 | eBioscience |
| CD44 | IM7 | v500 | 1 | eBioscience |
| CD44 | IM7 | eFluor450 | 1 | eBioscience |
| CD44 | IM7 | AF700 | 0.5 | BioLegend |
| CD45 | 30-F11 | PE-Cy7 | 1 | eBioscience |
| CD62L | MEL-14 | PE | 1 | BioLegend |
| CD62L | MEL-14 | BV711 | 1 | BioLegend |
| CD62L | MEL-14 | PE-Cy7 | 1 | eBioscience |
| CD69 | H1.2F3 | PE-Cy7 | 1 | eBioscience |
| CD80 | 16-10A1 | PerCP Cy5.5 | 1 | BioLegend |
| CD103 | PE | 2E7 | 1 | eBioscience |
| CD152 (CTLA-4) | UC10-4B9 | BV421 | 1 | BioLegend |
| CD278 (ICOS) | 15F9 | PE | 1 | BioLegend |
| CD278 (ICOS) | 7E.17G9 | BV421 | 1 | BD Bioscience |
| F4/80 | BM8 | PE | 1 | BioLegend |
| F4/80 | BM8 | APC | 1 | eBioscience |
| F4/80 | BM8 | PerCP-Cy5.5 | 1 | eBioscience |
| I-A/I-E (MHC-II) | M5/114.15.2 | PE-Cy7 | 1 | BioLegend |
| Foxp3 | MF-14 | AlexaFluor 647 | 5 | BioLegend |
| Foxp3 | MF-14 | PE | 2 | BioLegend |
| Foxp3 | FJK-165 | PerCP-Cy5.5 | 2 | BioLegend |
| Ki67 | SolA15 | PE-Cy7 | 1 | eBioscience |
| IFN-γ | 4S.B3 | PE-Cy7 | 1 | eBioscience |
| IFN-γ | XMG1.2 | PE-Cy7 | 1 | eBioscience |
| Biotin | Bio-18E7 | PE | 1 in 320 | Miltenyi |
| Biotin | Bio-18E7 | APC | 1 in 320 | Miltenyi |
| Viability | ef450; eF780 | None | 1:5000 | eBioscience |
| Armenian Hamster<br>IgG | HTK888 | biotin | 2.5 | BioLegend |
| Armenian Hamster<br>IgG | HTK888 | Pacific Blue | 0.24 | BioLegend |
| Armenian Hamster<br>IgG | HTK888 | AlexaFluor 647 | 0.24 | BioLegend |
| Rat IgG2a, $\kappa$ | eB149/10H5 | PE-Cy7 | 1 | BioLegend |
| Rat IgG1 | eBR41 | PE-Cy7 | 1 | eBioscience |
| Rat IgG2a, $\kappa$ | eBR2a | PerCP-Cy5.5 | 2 | BioLegend |
| Rat IgG2a, $\kappa$ | eBR41 | PerCP-Cy5.5 | 2 | BioLegend |
| Rat IgG2a, $\kappa$ | RTK4530 | AlexaFluor 647 | 5 | BioLegend |
| Rat IgG2a, κ | eBRa | AlexaFluor 647 | 1 | eBioscience |
| Rat IgG2a, κ | RTK4530 | PE | 2 | BioLegend |

### Western Blot

Lysis buffer was prepared by mixing 5 ml of 2X lysis buffer (100 mM HEPES, 300 mM NaCl, 20 mM NaF, 20 mM iodoacetamide) with 5ml ddH_2_O, and adding Complete Mini Protease Inhibitor Cocktail Tablet (Roche) and 100μl NP40 (BDH). Cells were suspended in 35 μl lysis buffer per 10^6^ cells and incubated on ice for 10 min. Samples were then centrifuged at 15000 rpm, 4°C for 10 min. Supernatant was aliquoted and 10 μl NuPAGE® Running Buffer (Life Technologies) added per 30 μl supernatant. 1M DTT reducing agent (Life Technologies) was then added at 1 in 50 to samples, which were heated at 70°C for 10 min. NuPage® 4-12% Bis-Tris Gels were loaded with sample and run in NuPage® SDS MOPS Running Buffer using the XCell SureLock™ Mini-Cell Electrophoresis System (all Life Technologies) for 50 min at 200 V, 210 mA. Meanwhile, blotting pads and filter paper were soaked in transfer buffer (1X NuPage® Transfer Buffer (Life Technologies), 10% methanol (VWR), and 1:1000 NuPage® Antioxidant (Life Technologies)). PVDF membranes (GE Healthcare) were soaked for 1 min in methanol, and rinsed in ddH2O. After running of the gel, blotting stacks were formed in the X Cell II™ Blot Module (Life Technologies) in accordance with manufacturer’s instructions. Proteins were transferred at 30 V for 1 h. Blots were blocked for 1 h at room temperature in 5% w/v milk (Marvel) in TBST (150 mM NaCl (AnalaR), 50 mM Tris-HCl (Melford), pH 7.6, plus 0.1% Tween 20).

After rinsing in TBST, blots were probed overnight at 4°C with anti-LC3 (Sigma) and anti-actin (Cell Signalling) in TBST with 5% w/v milk or BSA, respectively and 0.05% w/v NaN3. Blots were washed in TBST, then incubated for 1 h at room temperature with 1:25000 HRP-conjugated goat anti-rabbit IgG (Dako) or goat anti-mouse IgG (Dako) in 5% w/v milk in TBST. After further washes, blots were incubated at room temperature for 5 min in the dark with ECL detection reagent (GE Healthcare), made up in accordance with manufacturers’ instructions. Blots were transferred to a developing cassette, and film (Hyperfilm ECL, GE Healthcare) exposed for an optimum period. Film was developed using a Compact X4 developer (Xograph).

### Lipid kinase assay

5.1x10^6^ HEK293 cells were seeded in 10 cm dishes and transfected in duplicates the following day using Fugene HD at a 3:1 reagent:DNA ratio with the WT/Del21 constructs. After 24 h, cells were washed with cool PBS and lysed by scraping the wells with previously cooled CLB lysis buffer (1% TritonX100, 150 mM NaCl, 50 mM Tris pH7.4, 10% Glycerol, 1 mM CaCl2, 1 mM MgCl2, Protease/Phosphatase Inhibitors), and incubated on ice for 25 min. Lysates were spun at 15,000 x g for 15 min at 4°C and the supernatant fraction recovered. Protein concentration was determined by colorimetric assay (Bradford assay, Biorad). Protein G Dynabeads magnetic-agarose beads were used for immunoprecipitation of Myc-tagged proteins . The previous day, 20 μL slurry had been washed once with PBS before coupling overnight at 4°C with 3 μg of Myc antibody (Julian Downward’s) per sample.

After coupling, beads were washed once with CLB buffer, and incubated with 1mg of HEK293 cellular extract overnight at 4°C on a rotator. Samples were washed 3 times with ice-cold lysis buffer containing 0.1% Triton, and separated in two, one for kinase assay and the other one eluted in 50 μL of 1x sample buffer, which was boiled for 5 minutes to perform western blot. The kinase fraction was incubated with 25 μL of kinase buffer (20 mM Tris, pH 7, 67 mM NaCl, 10 mM MnCl2, 0.02% (w/v) NP40). Radioactive labelled γ-ATP (32P) (Hartmann SRP401) was used to quantify radioactive PtdIns3P by thin layer chromatography (TLC), as previously described (Chaussade et al., 2007). PI (Sigma P2517-5MG) was diluted in kinase buffer at a concentration of 1 μg/μL. 10 μL of PI per reaction was then added to 25 μL of IP. A master mix of 100 μM cold ATP per sample for 50 μL total volume reactions was prepared in kinase buffer. Once in the radioactive room, 0.1 μCi/μL of radiolabelled 32P-ATP per reaction was added to the master mix. 15 μL of the master mix was then added in a staggered manner to each sample and samples were incubated for 30 min at 37°C. Reactions were stopped by adding 100 μL of 1N HCL and inversion of the tubes. Once each reaction had been stopped, they were mixed vigorously for 2 min, after which 200 μL MeOH:CHCl3 (1:1) was added to each sample and then mixed again for a further minute. Samples were then centrifuged at 2,500 rpm for 5 min and the lipid phase was collected. 80 μL of MeOH: 0.1N HCl (1:1) was added to the samples and they were vortexed and spun at 2,500 rpm for 5 min. The lipid fraction was again collected and spotted onto a TLC plate, previously treated with 1% K-oxalate in MeOH:H2O 2:3 + 2 mM EDTA and ‘baked’ for more than 45 min at ∼124°C. Once the lipids were spotted, the plate was placed in a glass chamber containing 97.5 mL propanol-1, 46.5 mL water and 6 mL acetic acid (100%) for about 3 h. Prior to the experiment, the chamber had been previously stored in that buffer for a week. Once the TLC had run, the plate was dried in a fume hood, wrapped in cling film and placed on a phosphoscreen. The screen was analysed the following day using a Typhoon trio + phosphorimager (GE Healthcare Life Sciences).

### Polymerase chain reaction (PCR)

Deletion of exon 21 of the *Pik3c3* gene was confirmed by using the polymerase chain reaction (PCR). Treg cells were isolated from spleen and lymph nodes, enriched by immunomagnetic separation (EasySep™ Mouse CD4^+^ CD25^+^ Regulatory T Cell Isolation Kit II from StemCell) according to the manufacturer’s protocol. Fluorescence-activated cell sorting (FACS) was performed to sort for Treg cells (98% purity). Cells were lysed in 75 μl Cell Lysis Buffer for 30 min at 95°C, placed on ice for 10 min, and 75 μl Neutralization Buffer were added before the samples were snap-frozen.

The PCR reaction was performed in 50 μl, comprising 2 μl sample DNA and 48 μl reaction mixture containing per reaction 5 μl PCR buffer 10x (Invitrogen), 2 μl MgCl_2_ (50 mM), 1 μl dNTPs (10 mM), 1 μl of each primer (GCTGGTAGTACTGATGTTGC, GCATGGTCCTACTTTCTTCC and AGTCGAAGGTTGACTGTACC) with expected fragments as follows: 357bp (wt) and 455bp (*Pik3c3*^Δ21^).

### Quantification of mitochondrial DNA by quantitative PCR

Treg cells were isolated from spleen and lymph nodes, and lysed in the same way as described for the standard PCR procedure. Quantification of mtDNA copy number was performed using a Bio-Rad CFX machine, with 96-well plates (Bio-Rad Laboratories), Microseal ‘B’ plate sealers (Bio-Rad) and iTaq Universal Probes Supermix (Bio-Rad Laboratories). A final volume mastermix of 20 µl was prepared containing 1x iTaq Universal Probes Supermix, 300 nM forward and reverse primers, 200 nM Taqman probe and 10 ng of DNA sample. Actin was used as the reference (house keeping) gene. The qPCR reaction was amplified using the following conditions: initial denaturation at 95°C for 3 min, then 40 cycles of: denaturation at 95°C for 10 sec, followed by annealing and extension at 62.5°C for 1 min. Fluorescent levels were measured during each annealing and extension phase. The 2^-(ΔΔCt)^ method was used to calculate the relative abundance of mitochondrial DNA.

#### Primers

**Template for the standard curve:**

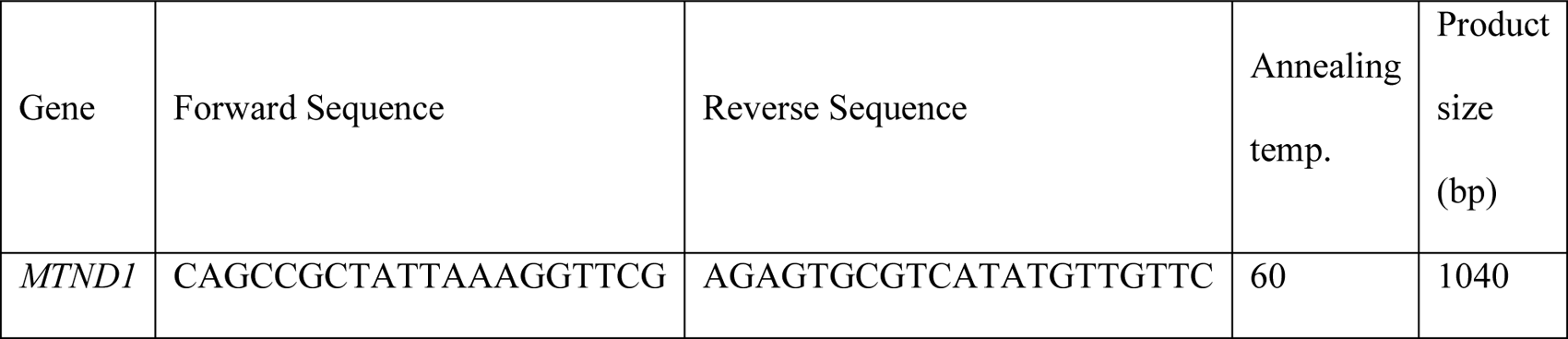

**qPCR:**

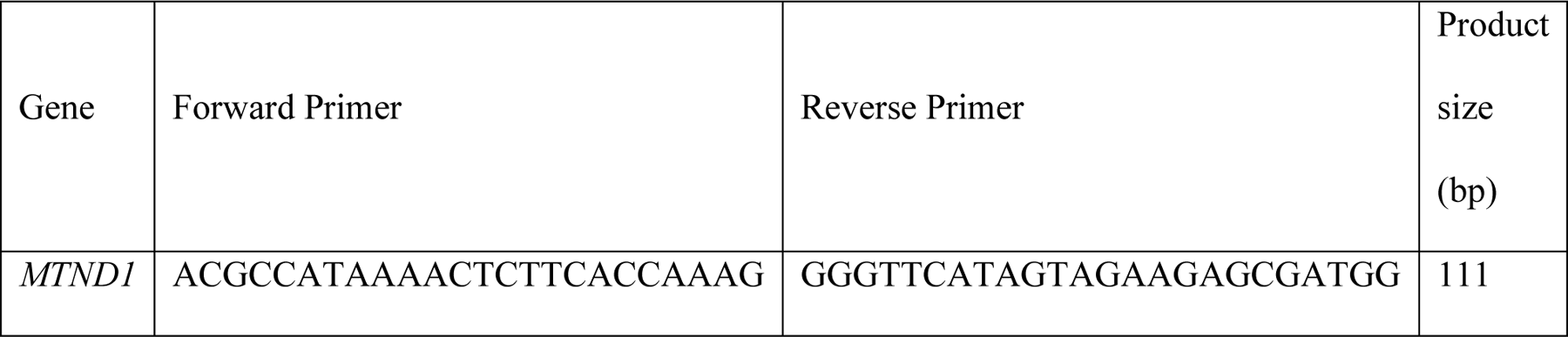

**Probe**:

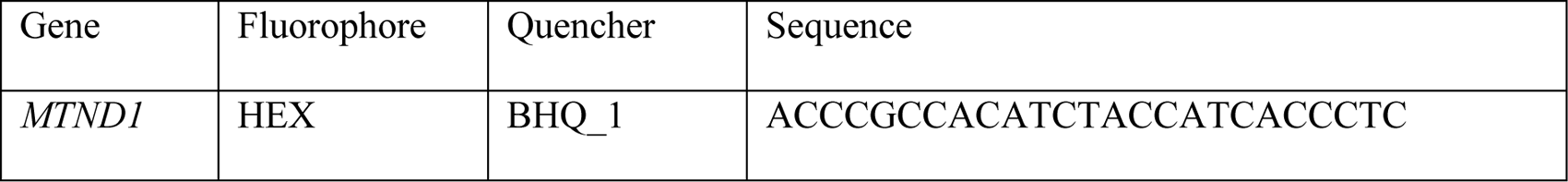

### Histology

For histopathological review, formalin-fixed tissues were sent to Propath UK, Hereford, for paraffin embedding, sectioning and haematoxylin and eosin (H&E) staining. Unstained slides were prepared for immunohistochemistry. To examine demyelination, samples embedded by Propath were sent to the Histopathology Service at the Royal Veterinary College (North Mymms, UK), for luxol fast blue staining.

### Seahorse XF Cell Energy Phenotype Test Assay

Prior to the assay, Agilent Seahorse XF96 Cell Culture Microplates were coated with Cell-Tak solution for 20 min at room temperature. On the day of the assay, Cell-Tak coated XF96 Cell Culture Microplates were allowed to warm to room temperature for 1 h. 0.5x10^5^ cells per well in 50 µl of assay medium (Seahorse XF Base Medium supplemented with 1 mM pyruvate, 2 mM glutamine, and 10 mM glucose, pH 7.4) were transferred to the XF96 Cell and centrifuged at 200*g* (zero braking) for 1 min. The plate was transferred to a 37°C incubator not supplemented with CO_2_ for 30 min; 130 µL warm assay medium were then gently added to each well, and the plate was returned to the incubator for 15–25 min. The sensor cartridge was loaded with the appropriate volumes described in the manufacturer’s protocol. The Seahorse XF Cell Energy Phenotype Test Assay was performed using 0.5 µM FCCP and 10µM oligomycin. At the end of the assay, protein levels were quantified to allow normalization. Supernatant was removed, cells washed once with PBS and lysed with 20 µl 0.2 M NaOH for 10 min. Subsequently, a BCA assay was performed according to the manufacturer’s protocol using 5 µl of the samples. After 30 min incubation at 37°C in a non-supplemented CO_2_ incubator, samples were read at 560 nm.

### Liquid chromatography mass spectrometry proteomics (LC-MS/MS) and data analysis

Treg cells from mosaic *Foxp3*^YFP-Cre/WT^*Pik3c3*^flox^ and *Foxp3*^YFP-Cre/WT^ control mice were isolated by magnetic enrichment followed by FACS, as described above. Sorted cells were washed twice in HBSS and snap-frozen in liquid nitrogen.

### Proteomics sample preparation and tandem mass tag (TMT) labeling

Cell pellets were lysed in 400 μl lysis buffer (4% sodium dodecyl sulfate, 50 mM tetraethylammonium bromide (pH 8.5) and 10 mM tris(2-carboxyethyl)phosphine hydrochloride). Lysates were boiled and sonicated with a BioRuptor (30 cycles: 30 s on and 30 s off) before alkylation with 20 mM iodoacetamide for 1 h at 22 °C in the dark. The lysates were subjected to the SP3 procedure for protein clean-up before elution into digest buffer (0.1% sodium dodecyl sulfate, 50 mM tetraethylammonium bromide (pH 8.5) and 1 mM CaCl2) and digested with LysC and Trypsin, each at a 1:50 (enzyme:protein) ratio. TMT labeling and peptide clean-up were performed according to the SP3 protocol. After labeling, samples were eluted into 2% DMSO in water, combined and dried in vacuo.

### Peptide fractionation

The TMT samples were fractionated using off-line high-pH reverse-phase chromatography: samples were loaded onto a 4.6 mm × 250 mm XbridgeTM BEH130 C18 column with 3.5 μm particles (Waters). Using a Dionex BioRS system, the samples were separated using a 25-min multistep gradient of solvents A (10 mM formate at pH 9 in 2% acetonitrile) and B (10 mM ammonium formate at pH 9 in 80% acetonitrile), at a flow rate of 1 ml min−1. Peptides were separated into 48 fractions, which were consolidated into 24 fractions. The fractions were subsequently dried, and the peptides were dissolved in 5% formic acid and analyzed by liquid chromatography–mass spectrometry.

### Liquid chromatography electrospray–tandem mass spectrometry analysis

For each fraction, 1 μg was analysed using an Orbitrap Fusion Tribrid mass spectrometer (Thermo Fisher Scientific) equipped with a Dionex ultra-high-pressure liquid chromatography system (RSLCnano). Reversed-phase liquid chromatography was performed using a Dionex RSLCnano high-performance liquid chromatography system (Thermo Fisher Scientific). Peptides were injected onto a 75 μm × 2 cm PepMap-C18 pre-column and resolved on a 75 μm × 50 cm RP C18 EASY-Spray temperature-controlled integrated column-emitter (Thermo Fisher Scientific) using a 4-h multistep gradient from 5% B to 35% B with a constant flow of 200 nl/min. The mobile phases were: 2% acetonitrile incorporating 0.1% formic acid (solvent A) and 80% acetonitrile incorporating 0.1% formic acid (solvent B). The spray was initiated by applying 2.5 kV to the EASY-Spray emitter, and the data were acquired under the control of Xcalibur software in a data-dependent mode using the top speed and 4 s duration per cycle. The survey scan was acquired in the Orbitrap covering the m/z range from 400–1,400 Thomson units (Th), with a mass resolution of 120,000 and an automatic gain control (AGC) target of 2.0 × 10^5^ ions. The most intense ions were selected for fragmentation using collision-induced dissociation in the ion trap with 30% collision-induced dissociation energy and an isolation window of 1.6 Th. The AGC target was set to 1.0 × 10^4^, with a maximum injection time of 70 ms and a dynamic exclusion of 80 s. During the MS3 analysis for more accurate TMT quantifications, ten fragment ions were co-isolated using synchronous precursor selection, a window of 2 Th and further fragmented using a higher-energy collisional dissociation energy of 55%. The fragments were then analyzed in the Orbitrap with a resolution of 60,000. The AGC target was set to 1.0 × 10^5^ and the maximum injection time was set to 300 ms.

### Processing and analysis of proteomics data

The data were processed, searched and quantified with the MaxQuant software package (version 1.5.8.3). Proteins and peptides were identified using the UniProt mouse database (SwissProt and Trembl) and the contaminants database integrated in MaxQuant, using the Andromeda search engine with the following search parameters: carbamidomethylation of cysteine, as well as TMT modification on peptide amino termini and lysine side chains, were fixed modifications; methionine oxidation and acetylation of amino termini of proteins were variable modifications. The false discovery rate was set to 1% for positive identification at the protein and peptide-to-spectrum match level. The dataset was filtered to remove proteins categorized as ‘contaminants’, ‘reverse’ and ‘only identified by site’. Copy numbers were calculated after allocating the summed MS1 intensities to the different experimental conditions according to their fractional MS3 reporter intensities. The accuracy of quantification was established using the following guidelines: proteins categorized as high accuracy had more than eight unique and razor peptides and a ratio for unique/unique + razor of ≥0.75; proteins categorized as medium accuracy had at least three unique and razor peptides, and a ratio for unique/unique + razor of ≥0.5; and any proteins below these thresholds were classified as low accuracy.

### Statistics and calculations

P values were calculated via a two-tailed, unequal-variance t-test on log-normalized data. Elements with P values < 0.05 were considered significant. Fold-change thresholds were established using a fold-change cut-off > 1.5 or < 0.67. The mass of individual proteins was estimated using the following formula: CN × MW/NA = protein mass (g cell−1), where CN is the protein copy number, MW is the protein molecular weight (in Da) and NA is Avogadro’s Constant. Pathway analyses were performed using the Database for Annotation, Visualisation and Integrated Discovery (DAVID) bioinformatics tools based on Kyoto Encyclopedia of Genes and Genomes (KEGG).

### Statistics and data analysis

Statistical analyses were carried out using GraphPad Prism (version 8.3.0). Where data were normally distributed, parametric tests were performed: unpaired students t-test with Welch’s correction was used when two groups were compared, or one-way ANOVA with Tukey post-test when three or more groups were compared. For data not following a normal distribution non-parametric tests were performed: a Mann-Whitney test was used when two groups were compared or the Kruskal-Wallis test with Dunns’ multiple comparison test was used when three or more groups were compared. Two-way ANOVA was used to compare mean differences in data sets influenced by the effect of two independent variables, such as in cases where different genotypes and drug treatment were involved. A two-tailed Student’s t-test was used for the statistical analysis of differences between two groups with Sidak’s correction when comparing two groups at multiple time points. Comparison of multiple groups was done using one-way ANOVA followed by Tukey’s *post-hoc* test. Significance is shown as * *p* < 0.05, ** *p* < 0.01, and *** *p* < 0.001.

## Acknowledgements

We thank Drs. Alexander Rudensky and Masaaki Komatsu for providing Foxp3^YFP-Cre^ and ATG7^flox^ mice, respectively. We thank Christina Rollings, Laura Spinelli and Doreen A. Cantrell for help and advise with mass spectrometry experiments. We thank members of the Babaraham Institute and University of Cambridge Animal facilities for their assistance and care for mice used in this study. We would also like to thank member of the Flow Cytometry facilities at the Babraham Instute and at the Department of Pathology, University of Cambridge, for expert assistance.

