## Supplemental figures 1 and 2 for "Lack of phosphatidylinositol 3-kinase VPS34 in regulatory T cells leads to a fatal lymphoproliferative disorder without affecting their development"

### Fig. S1. Gene targeting and characterization of the VPS34$^{Del21}$ mice

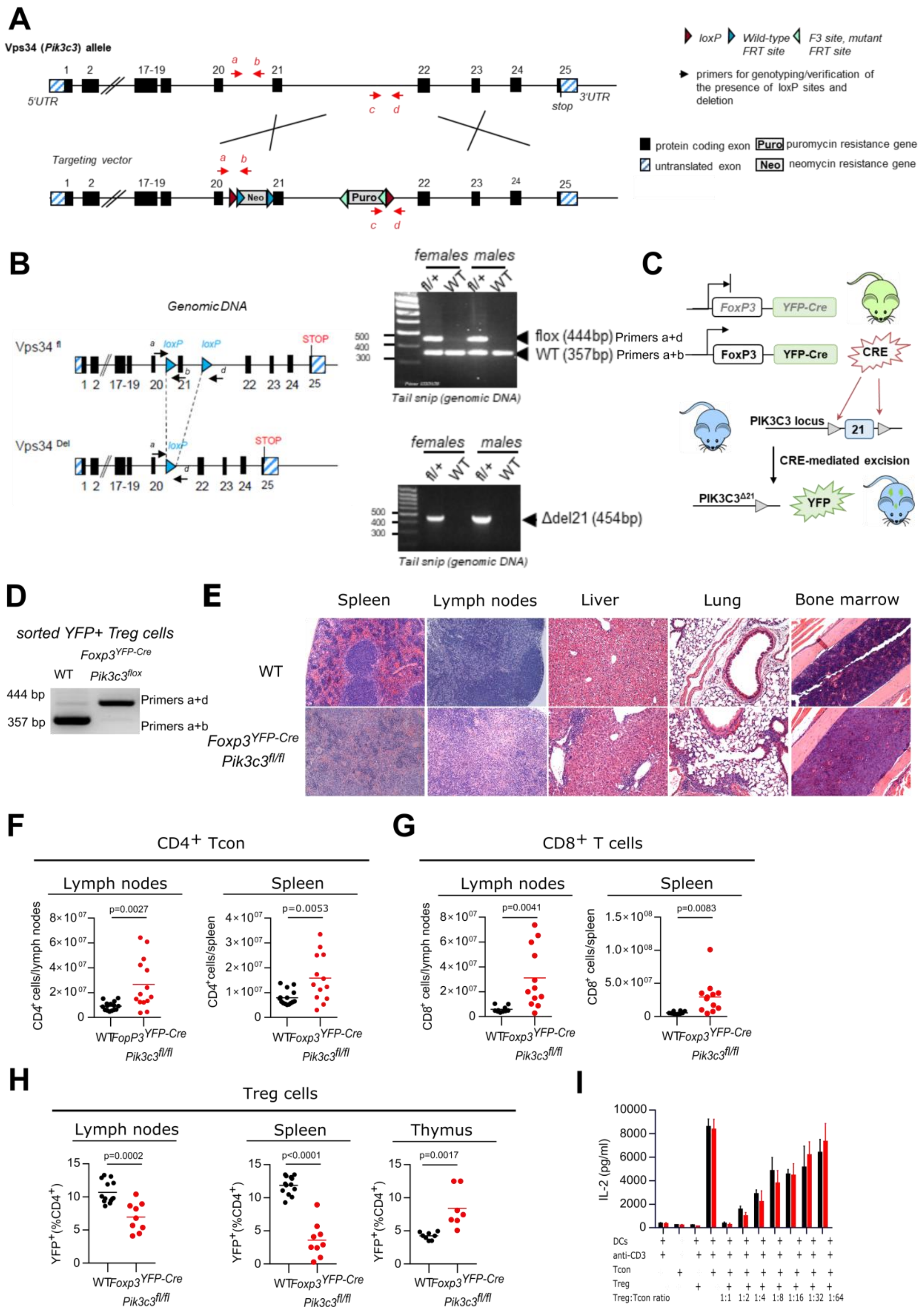

**A**) Gene targeting strategy to generate the conditional exon 21 deletion in the *Pik3c3* gene. The exon 21 encoding a critical stretch of the VPS34 kinase domain, was flanked by *lox*P sites, allowing an in-frame deletion upon exposure to Cre-recombinase. The FRT-flanked cassette encoding the *Pgk Neo* selection marker (Neo) was removed *in vivo* by breeding onto ACTB-Flp mice which express the FLP1 recombinase gene under the control of the *ACTB* (actin) promoter. **B**) *Pik3c3*$^{flox}$ mice were mated to Cre-deleter (B6.C-Tg(CMV-cre)1Cgn/J) transgenic mice which ubiquitously express Cre recombinase from the zygote stage of development (Schwenk et al., 1995). On the right, a representative image of agarose electrophoresis showing PCR products generated using the primer pair 1450_33,1450_34 (shown as “a”, “b”) for detection of heterozygous/homozygous WT and *loxP* alleles. For detection of deletion, the PCR primer pairs 1450_33 and 1451-38 (shown as “a” and “d” respectively) were used. Each PCR were performed on genomic DNA isolated from tail snips. **C**) Targeting strategy to generate Treg cell-specific VPS34-kinase dead mice where exon 21 of *Pik3*c3 (*Pik3c3*$^{flox}$) (Bilanges et al., 2017) is deleted specifically in Treg cells by the Cre-recombinase expressed under the control of the *Foxp3* locus. **D**) DNA from wild-type (WT) and VPS34-deficient Treg cells that were purified by FACS-sorting from 4 to 5-week-old from *Foxp3*$^{YFP\text{-}Cre}$ *Pik3c3*$^{WT}$ or *Foxp3*$^{YFP\text{-}Cre}$*Pik3c3*$^{flox}$ mice. A PCR product of 357 bp or 450 bp is detected for WT and VPS34-deficient Treg cells, respectively. **E**) Haematoxylin and eosin-stained sections from the spleen, inguinal lymph nodes, liver, lung, and femoral bone marrow of 32-day old Foxp*3*$^{YFP\text{-}Cre}$*Pik3c3*$^{flox}$ and wild-type *Foxp3*$^{YFP\text{-}Cre}$*Pik3c3*$^{WT}$ mice. 10x magnification. Absolute numbers of CD4$^+$ CD25$^-$ (**F**) and CD8$^+$ Tcon cells (**G**) in the lymph nodes (inguinal, brachial, and axillary) and spleen of Foxp*3*$^{YFP\text{-}Cre}$*Pik3c3*$^{flox}$ and wild-type *Foxp3*$^{YFP\text{-}Cre}$*Pik3c3*$^{WT}$ mice. **H**) Percentage of YFP$^+$ cells from CD4$^+$ cells in the lymph nodes (inguinal, brachial, and axillary), spleen, and thymus of Foxp*3*$^{YFP\text{-}Cre}$*Pik3c3*$^{flox}$ and *Foxp3*$^{YFP\text{-}Cre}$ *Pik3c3*$^{WT}$ control mice. Mice were between 4 and 5.5 weeks old. n = 5–15 mice per group. Statistical significance was determined using a two-tailed Student’s t-test. Results are pooled from 3 independent experiments. **I**) **I**) Tcon and Treg cells from *Foxp3*$^{YFP\text{-}Cre}$*Pik3c3*$^{flox}$ mice and control wild-type mice (WT) were co-cultured in the presence of dendritic cells and 1μg/ml anti-CD3 for 96 h. IL-2 in the supernatant was measured by ELISA.

**Fig. S2. *Foxp3*$^{YFP\text{-}Cre/WT}$*Pik3c3*$^{flox}$ mosaic mice are phenotypically healthy**

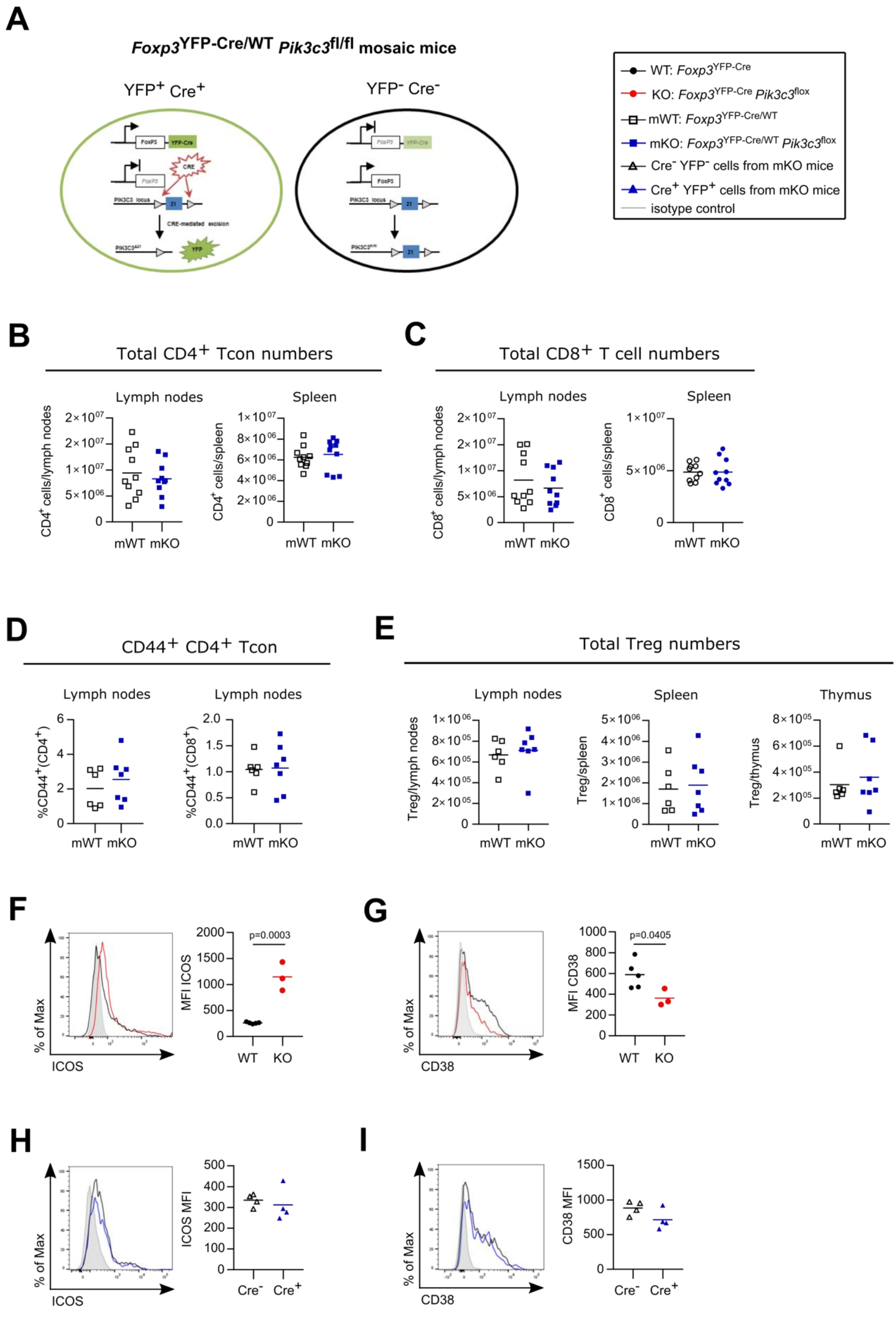

**A**) Mosaic knockout mice were generated by taking advantage of the localisation of the FoxP3 gene onthe X chromosome. Random X chromosome inactivation leads to the depletion of VPS34 in approximately 50% of Treg cells in female mice heterozygous for the FoxP3$^{\text{YFP-Cre}}$ transgene (FoxP3$^{\text{YFPCre/WT}}$). Accordingly, such mosaic female mice harbour two populations of Treg cells: a YFP$^{-}$ VPS34-sufficient and a YFP$^{+}$ VPS34-deficient Treg cell population. **B – C**) Absolute numbers of CD4$^{+}$ and CD8$^{+}$ T cells in spleen and lymph nodes (inguinal, brachial, and axillary) of Foxp*3*$^{\text{YFP-Cre/WT}}$*Pik3c3*$^{\text{flox}}$ mosaic (mKO) and wild-type *Foxp3*$^{\text{YFP-Cre/WT}}$*Pik3c3*$^{\text{WT}}$ (mWT) mice. **D**) Percentage of CD44$^{\text{high}}$ CD62$^{\text{low}}$ Tcon in the spleen and the lymph nodes of *Foxp3*$^{\text{YFP-Cre/WT}}$*Pik3c3*$^{\text{flox}}$ mosaic (mKO) and wild-type *Foxp3*$^{\text{YFP-Cre/WT}}$*Pik3c3*$^{\text{WT}}$ (mWT) mice. **E**) Absolute numbers of Treg cells in spleen, lymph nodes (inguinal, brachial, and axillary), and thymus of Foxp*3*$^{\text{YFP-Cre/WT}}$*Pik3c3*$^{\text{flox}}$ mosaic (mKO) and wild-type *Foxp3*$^{\text{YFP-Cre/WT}}$*Pik3c3*$^{\text{WT}}$ (mWT) mice.

**F – G** ) Mean fluorescence intensity (MFI) of ICOS (**F**) and CD38 (**G**) on splenic Treg cells of *Foxp3*$^{\text{YFP-Cre}}$*Pik3c3*$^{\text{flox}}$ mice (KO) and *Foxp3*$^{\text{YFP-Cre}}$ control mice (WT). **H – I** ) Mean fluorescence intensity (MFI) of ICOS (**F**) and CD38 (**G**) on splenic VPS34-deficient (Cre$^{+}$) and VPS34-sufficient (Cre$^{-}$) Treg cells from *Foxp3*$^{\text{YFP-Cre/WT}}$*Pik3c3*$^{\text{flox}}$ mosaic mice. *Foxp3*$^{\text{YFP-Cre}}$*Pik3c3*$^{\text{flox}}$ mice and the respective control mice were between 4 and 5.5 weeks of age. *Foxp3*$^{\text{YFP-Cre/WT}}$*Pik3c3*$^{\text{flox}}$ mosaic mice and the respective control mice were between 8 and 12 weeks of age. n = 3-7 mice per group. Statistical significance was determined using an unpaired two-tailed Student’s t-test (**B – G**), or paired two-tailed Student’s t-test (**H, I**). Results are pooled from 2 to 4 independent experiments.
